# Odor source localization in complex visual environments by fruit flies

**DOI:** 10.1101/179861

**Authors:** Nitesh Saxena, Dinesh Natesan, Sanjay P. Sane

## Abstract

Flying insects routinely forage in complex and cluttered sensory environments. Their search for a food or a pheromone source typically begins with a whiff of odor, which triggers a flight response, eventually bringing the insect in the vicinity of the odor source. The precise localization of an odor source, however, requires the use of both visual and olfactory modalities, aided by air currents that trap odor molecules into turbulent plumes, which the insects track. Here, we investigated odor tracking behavior in fruit flies (*Drosophila melanogaster)* presented with low- or high-contrast visual landmarks, which were either paired with or separate from an attractive odor cue. These experiments were conducted either in a gentle air stream which generated odor plumes, or in still air in which odor dissipates uniformly in all directions. The trajectories of the flies revealed several novel features of their odor-tracking behavior in addition to those that have been previously documented (e.g. cast-and-surge maneuvers). First, in both moving and still air, odor-seeking flies rely on the co-occurrence of visual landmarks with olfactory cues to guide them to putative odorant objects in the decisive phase before landing. Second, flies abruptly decelerate when they encounter an odor plume, and thereafter steer towards nearby visual objects that had no inherent salience in the absence of odor. This indicates that the interception of an attractive odor increases their salience to nearby high-contrast visual landmarks. Third, flies adopt distinct odor tracking strategies during flight in moving *vs.* still air. Whereas they weave in and out of plumes towards an odor source when airflow is present, their approach is more gradual and incremental in still air. Both strategies are robust and flexible, and can ensure that the flies reliably find the odor source under diverse visual and airflow environments. Our experiments also indicate the possibility of an olfactory “ working memory” that enables flies to continue their search even when the olfactory feedback is reduced or absent. Together, these results provide insights into how flies determine the precise location of an odor source.

## Introduction

Freely-flying insects live in a complex world that is both visually heterogeneous and odor-rich. This poses steep challenges in locating specific sources of odor which may include conspecific mates, food sources or oviposition sites. Moreover, these resources are often camouflaged in their natural surroundings or lack the distinctive visual features that identify them as putative odor sources. For insects flying in natural conditions, this task is confounded by the fact that flow conditions are turbulent and airflow can unpredictably change direction, which means that instantaneous odor signals may not provide reliable information about the location of an odorous object (Murlis, 1992; Vickers, 2000). Rather than diffusing along a smooth concentration gradient, odor signals in breezy conditions propagate as intermittent, filamentous plumes interspersed with clean air packets that greatly increase the range over which the odor molecules travel (Murlis, 1992; Willis et al., 1994). In hovering or slow-flying insects, these plumes are more laminar but disturbed by wing-induced upwind turbulence. This enhances odor sampling, but also alters spatial information about the location of an odor source (Sane and Jacobson, 2006).

It is well-known that airflow cues play a critical role in orienting flying insects toward an odor source (e.g. Kennedy and Marsh, 1974). Airflow collimates the odor cues thereby providing important directional cues to the odor-tracking insect. For instance, during active plume-tracking in laminar airflow, insects fly upwind aligning with the odor plume. Such behavior typically consists of two key aerial maneuvers. First, upon encountering an odor plume, insects perform *surging* maneuvers, which involve flying forward in the direction of the upwind odor source. However, if they lose track of the plume, they *cast* orthogonally to the plume axis to regain contact with the odor packets. The combination of casting-and-surging naturally channels the insect towards the source of odor (Farkas and Shorey, 1972; Kennedy, 1983; Vickers and Baker, 1994; Vickers, 2000).

For the *cast-and-surge* strategy to be effective, airflow must be relatively uniform and laminar. However, under most natural conditions, airflow can be quite erratic which means insects require supplementary information from other sensory modalities, especially vision. For example, fruit flies rely on wide-field visual cues during odor tracking (Frye et al., 2003; Budick and Dickinson, 2006; Duistermars and Frye, 2008), and moths utilize ambient visual cues to estimate airflow direction (e.g. Kennedy and Marsh, 1974). The sensing and processing by one sensory modality is often influenced by feedback from another modality during active behaviors. In odor tracking fruit flies, the presence of odor cues can modify optomotor responses thus enhancing their chances of honing in on visual features while maintaining a constant heading (Chow and Frye, 2008).

As they approach an odor source, insects rely on local visual cues to find odor objects (Raguso and Willis, 2002). Indeed, visual landmarks become attractive to insects *only* if odor is present (e.g. fruit flies, van Breugel and Dickinson, 2014; mosquitoes, van Breugel and Dickinson et al., 2015). However, if visual cues are indistinct or ambiguous, flies may increase their reliance on odor cues to find the sources of odor (e.g. in the Tephritid apple fly *Rhagoletis pomonella* (Walsh); Aluja and Prokopy, 1993). Under tethered conditions, local visual landmarks by themselves appear insufficient to help orient flies toward odor plumes, and may require wide-field visual cues such as panoramic background from the surroundings to navigate to the odor source (Duistermars and Frye, 2008).

Although the above studies demonstrate the importance of combining olfactory, airflow and visual cues in guiding insects to the general vicinity of an odor source, they do not reveal how they pinpoint the precise location of an odor source within complex visual environments. Here, we conducted experiments to test the hypothesis that local landmarks are essential in guiding flies to an odor source in the final stages before making a decision to land. Such a strategy is especially necessary in the relative absence of ambient airflow in which odor gradient tracking followed by guidance *via* local landmarks can provide an additional reference to locate odor objects. Our broad approach involved presenting simultaneous but spatially-staggered visual and odor cues to compel the flies to choose between them, in both the presence and absence of laminar airflow. Using high-speed videographic reconstructions of their 3D flight trajectories at high spatial resolution, we deconstructed how the combined odor and visual stimuli influenced the trajectories of flies prior to landing. Our data provide few simple rules used by flies to pinpoint the location of an odor source.

## Materials and Methods

We used 2-3 days old Canton-S flies from a culture maintained at the National Centre for Biological Sciences campus in Bangalore. Fly stocks were maintained at room temperature between 24-27°C, and in a 12 hr: 12 hr light-dark cycle. Prior to the experimental trials, the flies were starved overnight for ∼12 hours to increase their motivation for foraging. They were provided with water soaked paper during starvation period to prevent dehydration. Experiments were conducted during flies’ photoperiod to ensure robust flight activity.

*Visual cues*: We used two objects with different visual contrast that acted as low- or high-contrast visual landmarks for the flies.

*Low-contrast landmark*: A transparent glass capillary (length = 100 mm, diameter = 1 mm) placed within a small Plexiglas® holder tipped with cotton ball constituted the low-contrast visual landmark.

*High-contrast landmark*: We threaded a black spherical bead (diameter = 6 mm) on the glass capillary described above. The bead subtended an angle of ∼5° on the fly retina at a distance of ∼80 mm from the bead (our region of interest), and constituted a high-contrast local visual landmark for the approaching flies.

*Odorous landmarks*: Odor cues consisted of 10 μl of apple cider vinegar (5% vinegar syrup, Zeta Food Products, Stockholm), placed on the black bead (*high-contrast odorous landmark*) or cotton tip of the capillary (*low-contrast odorous landmark*) depending on the experimental treatment.

*Wind tunnel and filming apparatus*: We used a custom-made, calibrated low-flow wind tunnel to generate laminar airflow for experiments conducted in the presence of airflow. Flies were released in the test chamber (1200 mm X 280 mm X 280 mm), within which odor and visual cues were placed (Fig. 1A). For experiments involving the presence of airflow, the value of laminar airflow within the test section was set to 0.1 m/s, which is within the range of naturally-occurring airflow values. Air speed was measured using a hot-wire anemometer (Kurz, 490-IS portable anemometer, Monterey, California, USA for details see Sane and Jacobson, 2006). For experiments conducted in still air, wind tunnel motor was switched off and both ends of the wind tunnel were sealed using Plexiglas® sheets, thereby reducing ambient flow to values that were too low to be measured by the anemometer. The odorous landmark was then placed within the still chamber and odor was allowed to diffuse for ∼20 minutes. After this, flies were released inside the wind tunnel. To enhance the contrast between fly and the background, we lined the sides and base of the wind tunnel working section with white paper, which was then backlit by four 50 W halogen lamps to provide illumination for flies to track visual objects. A 150-watt metal halide lamp on top of the wind tunnel provided sufficient illumination for high-speed filming. We placed an additional red filter on the 150 W lamp to ensure that we used illumination of only wavelengths above 610 nm. Because fruit flies are relatively insensitive to light of wavelengths above 600 nm (Heisenberg and Wolf, 1984), we chose illumination of wavelengths above 610 nm for minimal impact on the flight behavior. The average illumination within the chamber was ∼350 lux, measured using a light meter (Center 337, Center Technology Corporation, Taipei, Taiwan).

**Figure 1:**
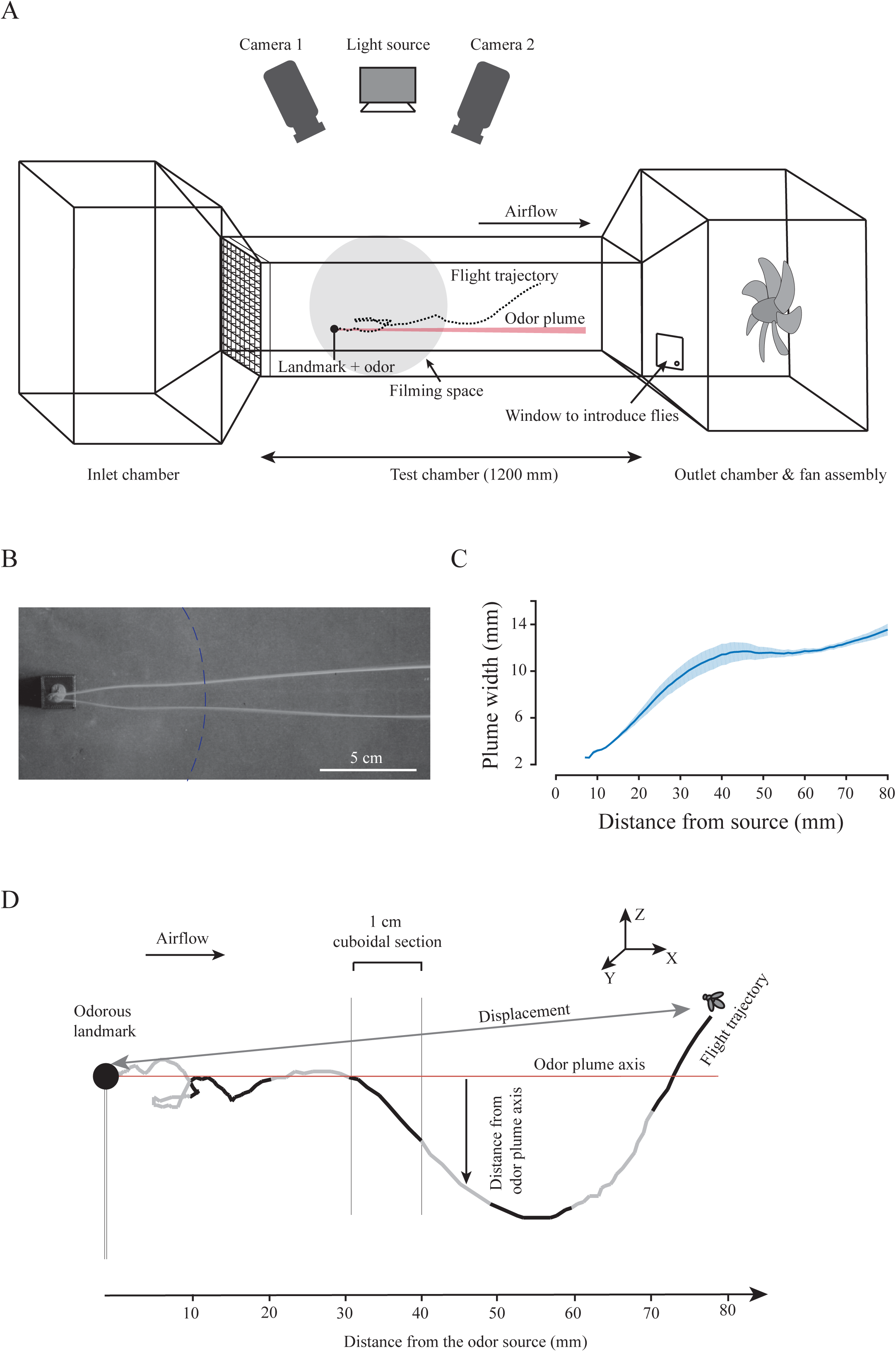
Experimental setup and flight variables. (A) Flies tracked the odor plume inside a customized wind tunnel of test chamber dimensions 1200 mm X 280 mm X 280 mm. The flies tracking the odor plume (red band) were filmed at 100 fps using two high-speed cameras mounted above the wind tunnel (approximate filmed region shown as a grey shaded circle) and their 3D flight trajectories could be reconstructed from these images. (B) A raw image of a laminar smoke plume from a low-contrast landmark source (view from above, see methods). The dashed line shows the 80 mm radial cut-off used in our experiments. (C) Change in plume width vs. distance from the source over 80 mm distance. The dark blue line shows the mean plume width and the light blue band shows the standard error around the mean (N = 4). (D) Schematic of a fly's typical approach to an odor source. The trajectories are broken into black and grey lines, each depicting flight along 10 mm stretches. The odor plume axis (red line) indicates the alignment of the odor plume, determined using a photo-ionization detector. We calculated speed, flight duration, tortuosity and hover duration to quantify the flight behavior in a spherical region of 80 mm diameter from the odor source (see *Methods*).

We filmed the flight trajectories at 100 frames per second using two high-speed cameras (Phantom v7.3 / Miro eX4, Vision Research, Wayne, New Jersey, USA) placed above the wind tunnel. We introduced approximately 5 or 6 flies inside the wind tunnel in every trial to reduce the waiting time for observing a landing event. Typically, only one fly approached the odor source at any given time and after first fly landed on any landmark, the trial was terminated. In rare cases, if multiple flies approached the odor source simultaneously, such trajectories were excluded to avoid the confounding effects of competitive social interactions between flies. Before starting another trial with a new set of flies, we flushed out flies from the previous trial from the wind tunnel to ensure that we recorded only the innate responses of naïve flies. One flight trajectory was filmed in each trial and 3D trajectories within 80 mm distance range from the odor source were analyzed.

### Treatments

We tested the responses of flies towards different arrangements of odor and visual cues. Each specific set of odor and visual cue arrangement constituted a *treatment.* All treatments and corresponding results are represented using icons and summarized in Table 1.

**Table 1:**
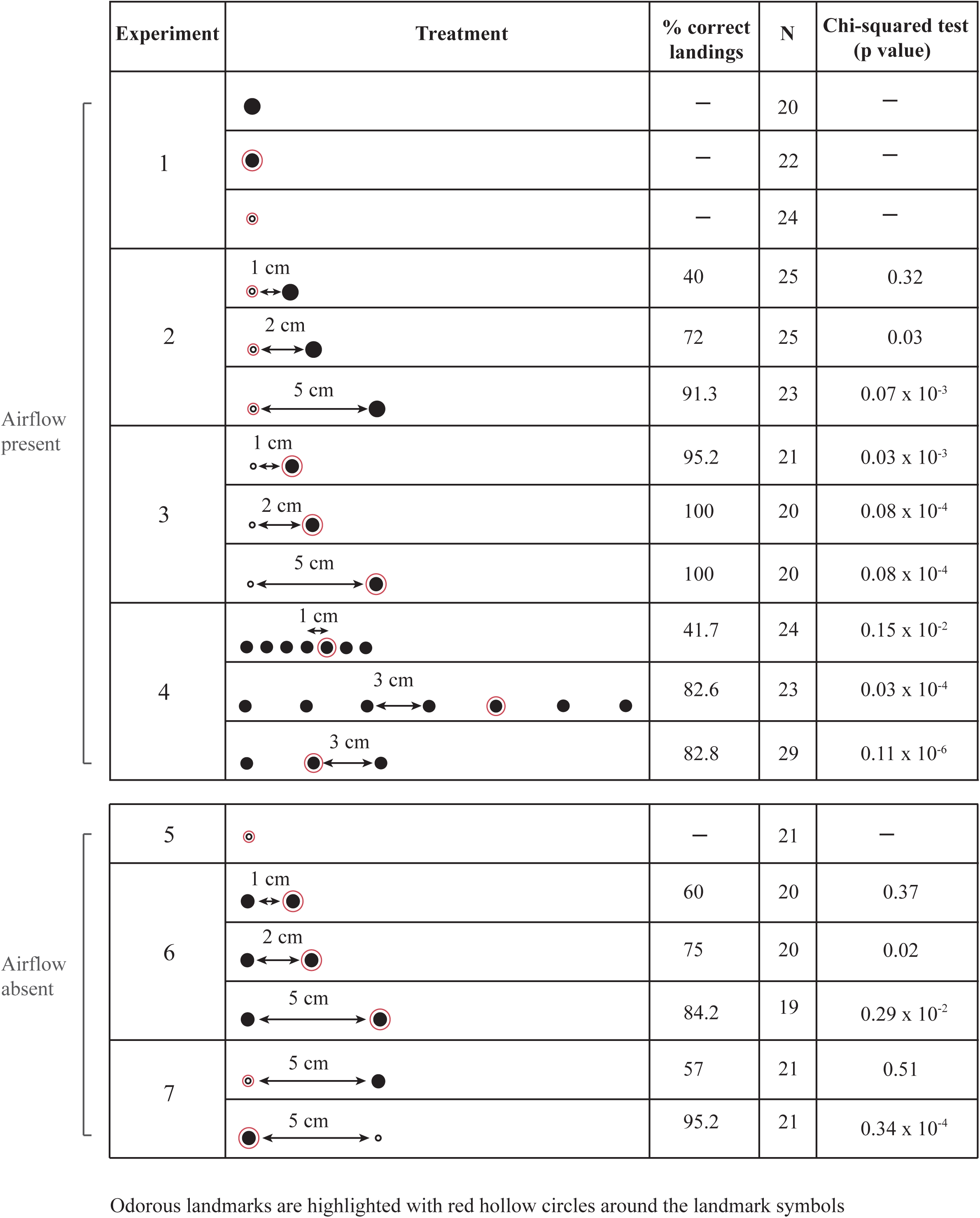
A summary of experiments. Small open circles denote low-contrast landmark, and larger solid circles denote high-contrast visual landmarks. A concentric red circle around the landmark represents the presence of odor. First 9 rows represent experiments in presence of airflow (and odor plume), whereas the bottom 5 represent experiments in still air. Correct landings are defined as the landings on the odorous landmark. The sample sizes (N) per treatment and p-value of Chi-squared test for each experiment are shown in 4^th^ and 5^th^ columns. The Chi-squared test compares the observed landing frequency with the expected frequency due to random landings on available landmarks (p < 0.05 indicate non-random landings).

### Experimental design

In all, we conducted seven experiments in which naïve flies were required to identify the odor source in presence of visual landmarks. In the first set, the flies flew in the presence of a constant 0.1 m/s airflow (experiments 1-4) whereas a second set was conducted in still air (experiments 5-7). We systematically varied the arrangement of visual landmarks around a single odorous landmark. Each experiment contained multiple treatments, with a fixed combination of odor, visual and airflow conditions for which individual responses of flies were filmed over multiple trials. These experiments are described below.

### Presence of airflow

*Experiment 1: Responses to individual odor and visual cues, and their combination:*

Flies were flown under three conditions:

1. A single *high-contrast non-odorous landmark,* to observe the innate responses of flies towards a visual cue in absence of odor cues.
2. A single *high-contrast odorous landmark,* to observe flight responses towards combined visual and odor cues and,
3. A single *low-contrast odorous landmark,* to observe flight responses towards odor cues with the low-contrast visual cue, which was a glass capillary with cotton tip to facilitate landing of flies.

*Experiment 2: Responses to decoupling of odor and visual cues:* We decoupled the odor and visual cues such that the *high-contrast non-odorous landmark* was kept separate from the *low-contrast odorous landmark* by a distance of 1, 2 and 5 cm respectively. These three treatments were compared with single *low-contrast odorous landmark* from experiment 1.

*Experiment 3: Control:* As the control case for above experiment, we switched the positions of odorous and non-odorous landmarks, now keeping the *high-contrast odorous landmark* and *low-contrast non-odorous landmark* separated by 1, 2, and 5 cm respectively. These three treatments were compared with single *high-contrast odorous landmark* from experiment 1.

*Experiment 4: Odor source localization in visual clutter:* To determine how flies identify odor sources within visually cluttered environments, we varied the number and density of landmarks around the odor source.

Flies were flown under three conditions of visual clutter:

1. *High-density visual clutter of seven objects:* We arranged seven high-contrast landmarks each separated by 1 cm in a single row. Odor was contained in an off-center (fifth) landmark, to ensure that flies landing on objects was not an indirect consequence of the natural centering response displayed by many insects when flying through confined spaces (e.g. Srinivasan et al., 1996). Our experimental design forced the fly to actively break symmetry to find odor source, thereby avoiding bias toward the central landmark.
2. *Low-density visual clutter of seven objects:* Seven high-contrast landmarks were each separated by 3 cm with the fifth landmark from right (approaching upwind) containing odor;
3. *Low-density low visual clutter of three objects*. Three high-contrast landmarks were arranged in a row, each separated from its nearest neighbor by 3 cm. Although the middle landmark was the odor object here, it was off-center.

### Absence of airflow

Airflow breaks the directional symmetry, and insects typically respond by flying in the upwind direction during odor tracking. In the absence of airflow, odor spreads primarily by diffusion in all directions. How do flies resolve the challenge of finding odor source in still air? We created the conditions required to address this question in Experiments 5-7.

*Experiment 5:* A single *low-contrast odorous landmark* was presented to the flies flying in the wind-tunnel with the fan switched off and wind tunnel sealed as previously described.

*Experiment 6:* We tested if flies were capable of distinguishing between two identical high-contrast landmarks of which only one was odorous, placed 1, 2 and 5 cm apart respectively.

*Experiment 7:* To determine the effect of visual contrast on odor tracking behavior in the absence of airflow, we designed two treatments. In one treatment, we placed a *low-contrast odorous landmark* separated by a *high-contrast non-odorous landmark* by 5 cm. As a control treatment, we placed *high-contrast odorous landmark* separated by a *low-contrast non-odorous landmark* by 5 cm.

The sample sizes and landing preferences of flies in the above treatments are provided in Table 1. These experiments allowed us to make systematic observations of landing preferences and perform the analysis of flight trajectories to explore the behavioral rules underlying odor tracking in fruit flies.

### Quantification of airflow

*Laminarity:* To ascertain the laminarity of the wind tunnel, we used two methods. First, we placed a hot-wire anemometer at separate points within the test section and verified that the value of airspeed at various points in space was identical, and at each point it held a constant value in time (see also Roy Khurana and Sane, 2016). Next, we determined laminarity for the airflow conditions for various treatments, to ensure that the presence of objects within the wind tunnel did not introduce turbulence in the internal flows. From a theoretical perspective, such turbulence is unlikely for reasons outlined below. For an object of characteristic length L placed in a fluid with velocity ***V*** and kinematic viscosity *v*, the Reynolds number (*Re*)is given by:

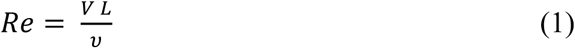

For airflow of 0.1 m/s, kinematic viscosity of 1.57 x 10^-5^ m^2^/s (dry air at 300 K) and characteristic lengths from 1 mm (single capillary) to 10 mm (smallest separation distance between two landmarks), the Reynolds numbers range from ∼ 7-70, well within the laminar regime. We tested this expectation using flow visualization in the wind tunnel (see Supplementary video 1).

### Plume Visualization

To visualize the odor plume, we seeded the flow with smoke generated using moldable incense clay, mimicking the odor source in our experiments. We simulated the following treatments, which encompassed the odor plume conditions in all experiments:

1. Capillary (i.e. low-contrast object).
2. Capillary generating smoke, with a spherical bead (6 mm diameter, i.e. high-contrast object) at 1 cm.
3. Capillary generating smoke, with a spherical bead at 2 cm.
4. Spherical bead.
5. Spherical bead generating smoke, flanked by two beads at 1 cm.

We filmed the smoke plume at 24 fps using a calibrated high-resolution high-speed camera (Phantom VEO 640L, Vision Research, Wayne, New Jersey, USA) which directly viewed the object from above. The wind tunnel was set at 0.1 m/s. For every smoke plume treatment, we filmed four trials saving a minimum of 100 frames per video, which were processed using Fiji (Schindelin et al., 2012) software. We recursively subtracted the background from each image in the stack to obtain the averaged steady-state image of the axisymmetric plume. Undetected gaps in the plume were interpolated using piecewise Cubic Hermite spline. By adjusting the threshold and filtering this image with a median filter to remove salt-and-pepper noise, we obtained a binary form of this image, which was imported into MATLAB and digitized using a custom code (Supplementary Figure 1A-E). We obtained the plume width with respect to the source distance by pooling the data at a resolution of 1 mm (Supplementary figure 1-F). The plume width of the smoke plume saturated to become roughly cylindrical about 4-8 cm from the odor source. The presence of neighboring spherical beads only slightly affected the plume width, causing it to vary between 1-1.6 cm in diameter. For all calculations relating to odor encounters, we set 1.6 cm as the diameter of the plume.

### Data Acquisition and Analysis

Two cameras simultaneously recorded the fly’s trajectory as it approached the odor source. The fly’s position was digitally marked in each camera view and their 3D position reconstructed using custom MATLAB software (Hedrick, 2008). The extracted trajectories were processed through a 4^th^ order Butterworth filter with a cut-off of 30 Hz. The *high-contrast landmark* used in this study subtended an angle of ∼5° at ∼8 cm distance, ensuring that this angle was slightly greater than the smallest inter-ommatidial angles of ∼4.5° in *Drosophila* (Gonzalez-Bellido et al, 2011). We digitized and analyzed only the flight trajectories within the 8 cm radius from the odor source. From the 3D flight trajectories, we calculated several flight variables of which four best captured the spatio-temporal features of their trajectories (Fig 1B):

1. *Flight speed*: the average speed of a fly.
2. *Flight duration*: the total duration of flight trajectories.
3. *Hover duration:* the total duration spent by a fly at speeds less than 37.5 mm/s (hover speed). We chose this cut-off speed because it represents a value closer to true hover (assuming a body length of 3 mm, which is less than 5 % body length traversed over a single wing beat duration of ∼4 ms).
4. *Tortuosity:* ratio of total distance travelled by the fly to its displacement.

Flight activity was non-uniform near the odor source due to steady deceleration of flies as they narrowed their search. These changes depend on the distance of flies from the odor source. Hence, we segmented the volume in front of the odor source into 784 cm^3^ (1 cm X 28 cm X 28 cm) cuboids along the length of the wind tunnel (Fig 1B). For each treatment, we separately analyzed the free-flight behavior in each spatial zone and statistically compared changes in flight variables across these segments. The calculated values of flight trajectory variables were not normally distributed (Lilliefors test for normality at p<0.05) and did not have equal variances (Bartlett test for equal variance at p<0.05). Hence, we used non-parametric tests to compare the statistical significance of observed differences in the flight variable values. To detect whether any groups were statistically different (at p<0.05) from the other groups, we used the Kruskal-Wallis test, a non-parametric version of ANOVA. If this test indicated significant differences between one or more groups, we used the post-hoc Nemenyi test to compare each group in a pairwise manner and identified which specific treatments were different from each other.

## Results

### The presence of odor cues alters the response of flies toward visual landmarks

When presented with a *high-contrast non-odorous landmark*, flies maintained an upwind heading but did not approach the visual landmark (Fig 2A). However, the landmark became attractive to flies when it emitted an appetitive odor (Fig 2B, C). Before landing, flies aligned themselves along the plume axis as they approached the *high-contrast odorous landmark* (Supp. Fig 2B, C), whereas their flight towards the non-odorous landmark was not directed along any specific axis (Supp. Fig 2A). Flies also flew at significantly slower speeds (Fig. 2D) and for longer duration (Fig. 2E), and their trajectories were more tortuous (Fig. 2F) in presence of odor cues. In addition, the hover duration was also significantly greater in the vicinity of an *odorous landmark* than the *non-odorous landmark* (Fig 2G). Thus, the presence of odor increased flight activity in general. Flight trajectories of flies approaching high- and low-contrast odorous landmarks were not statistically different from each other (Fig 2D-G). This shows that the presence of odor cues was necessary and sufficient for flies to seek out a visual landmark, even when it was of a lower contrast.

**Figure 2:**
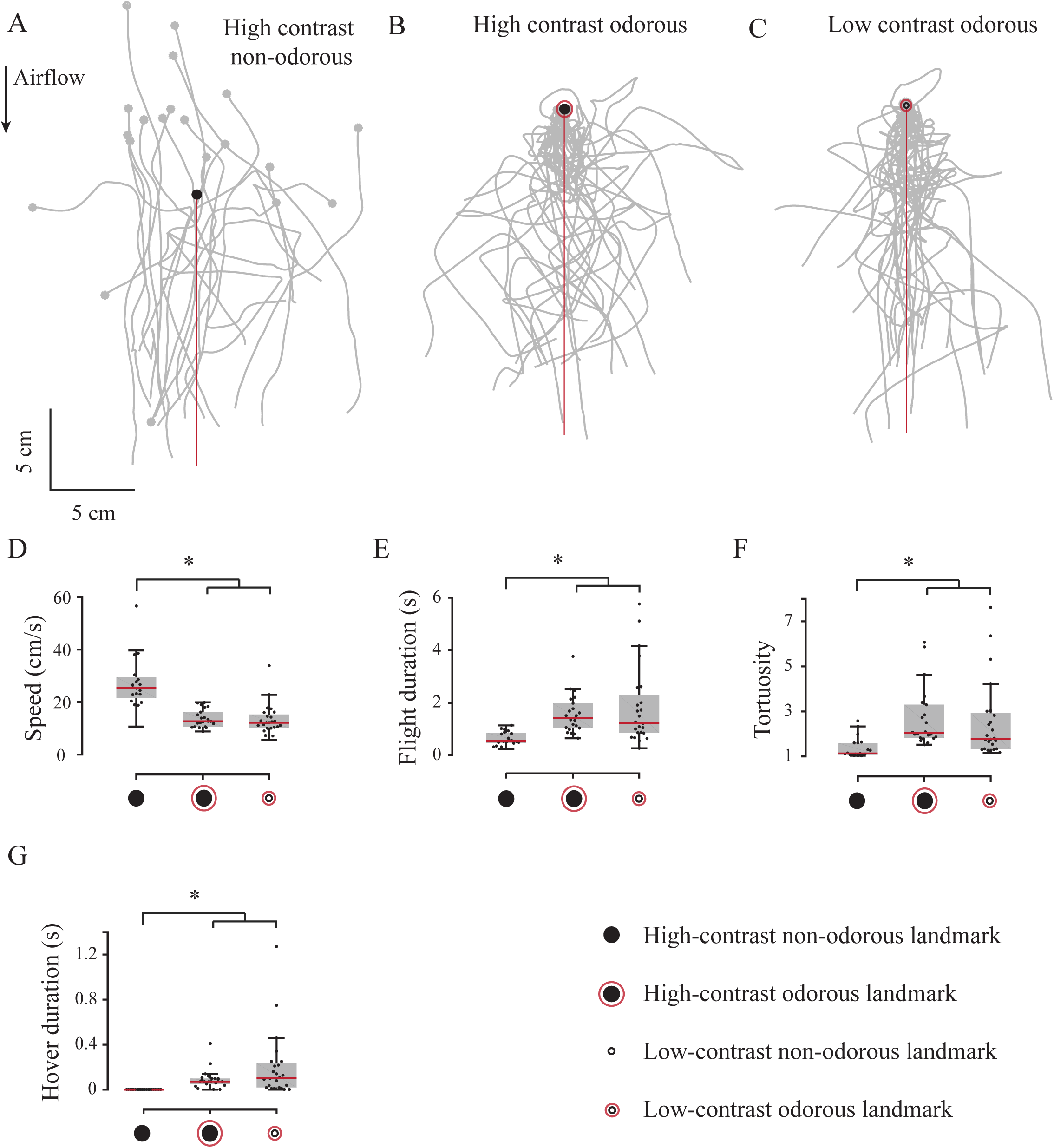
Flight behavior in the presence of odorous and non-odorous landmarks. Flight trajectories (grey) in the presence of (A) a high-contrast non-odorous landmark (N=20), (B) a high-contrast odorous landmark (N=22), and (C) a low-contrast odorous landmark (N=24) respectively. The flies flew towards an odor source that was either high-contrast (filled circle) or low contrast (open circle) by tracking an odor plume along its axis (red line). Odorous landmarks are depicted by a concentric red circle around the circles depicting visual objects. We compared between these treatments for average speed (D), the total flight duration (E), tortuosity (F) and the hover duration (G) of the flies, depicted as box-and-whisker plots. The height of the box indicates the range of central 50 % of data around the median (red line). The length of whiskers represents data that is within 1.5 times the interquartile range. Outlier data lie outside the whiskers, but are included in analysis. Asterisks represent statistically significant comparisons (p<0.05, Kruskal Wallis test, Nemenyi test) in all figures. The conventions of depicting odorous objects, box plots and statistical tests is followed throughout this manuscript.

### Flies integrate odor and visual cues prior to landing

We next presented flies with two choices for landing – a *low-contrast landmark* and a *high-contrast landmark*, of which only one was odorous. The two landmarks were separated by 1, 2 or 5 cm respectively in separate treatments. In the first set of experiments, the odor was paired with a low-contrast landmark (Fig 3 A-E), and in a second set, with a high-contrast landmark (Fig 4 A-E). If the presence of odor cues is sufficient to determine the landing site, then the landings should occur only on the odorous landmark regardless of the presence of a nearby landmark. However, flies showed some likelihood of landing on the *high-contrast non-odorous landmark* rather than the *low-contrast odorous landmark* (Fig 3A-C; Table 1), with the frequency of incorrect landings gradually decreasing as separation between the two objects increased (Fig. 3A-C, upper panels; Table 1). In contrast, when given a choice between *high-contrast odorous landmark vs. low-contrast non-odorous landmark*, flies always chose the former (with a sole exception, Fig. 4A), regardless of the separation distance between landmarks (Fig 4 A-C, Table 1). Thus, the co-occurrence of odor cue with a single high-contrast visual cue is sufficient to guarantee that flies will land on that object.

**Figure 3:**
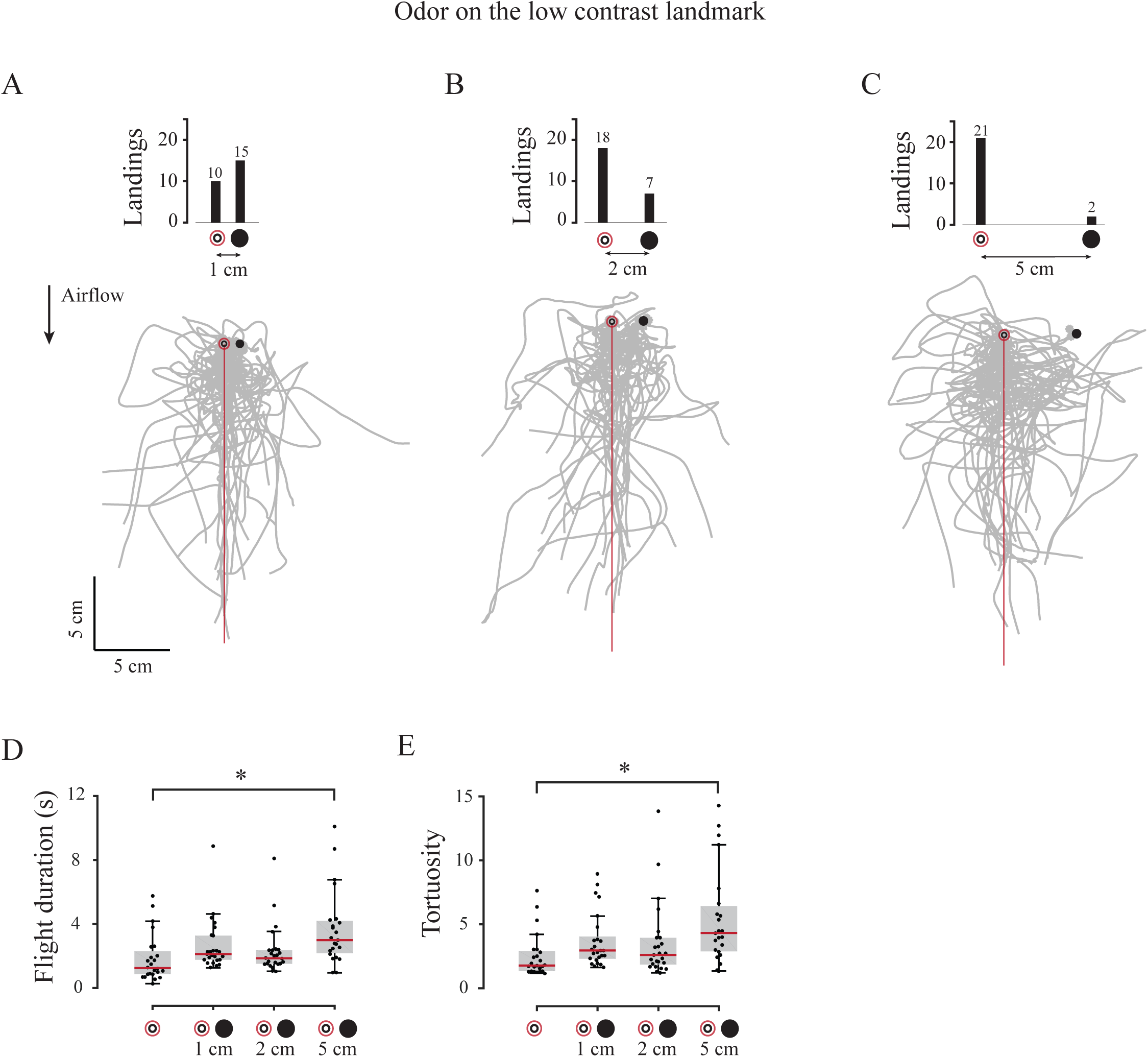
Landing preference and flight behavior in the presence of segregated odor and visual cues. Flight trajectories (grey) in the presence of an odorous low-contrast landmark separated from a non-odorous high contrast landmark by (A) 1 cm (N = 25), (B) 2 cm (N = 25) and (C) 5 cm (N = 23), respectively. Here and elsewhere, the bar plots above the trajectories, and the associated numbers indicate the absolute number of landings on each landmark. Also plotted are (D) flight duration and (E) tortuosity of the flies prior to landing.

**Figure 4:**
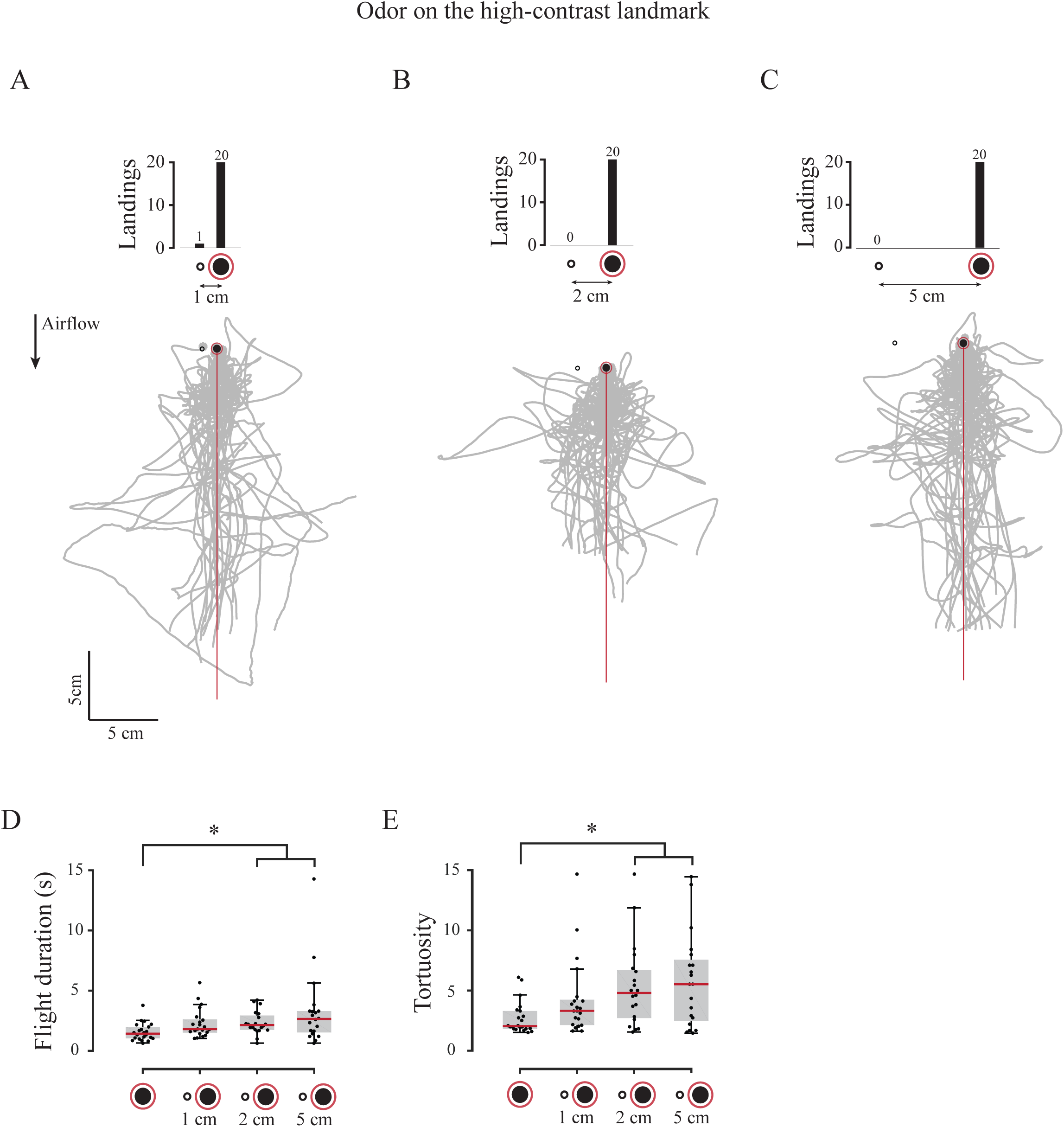
Landing preference and flight behavior on a high-contrast odorous landmark separated from a low-contrast non-odorous landmark. Flight trajectories (grey) in the presence of a high-contrast odorous landmark and a low-contrast non-odorous landmark at (A) 1 cm (N = 21), (B) 2 cm (N = 20) and (C) 5 cm (N = 20) separation respectively. As before, bars above the plots indicate the landing preferences on each landmark. The presence of a low-contrast non-odorous landmark near the high-contrast odorous landmark significantly increased both their (D) flight duration and (E) tortuosity prior to landing.

The flight duration and tortuosity values for flies approaching a *high-contrast non-odorous landmark* placed at 5 cm from a *low-contrast odorous landmark* were significantly different from flies approaching a *low-contrast odorous landmark* (Fig. 3 D-E), but the flight parameters were largely similar for smaller separations. Thus, it costs the flies some time to investigate the *high-contrast landmark* when it is not the source of odor which means that their search strategy is influenced by neighboring visual landmarks. The presence of a *low-contrast non-odorous object* also affects the trajectories of the flies if it is placed near a *high-contrast odorous object*, and flight duration and tortuosity were significantly greater when these were 2 and 5 cm apart (Fig. 4D, E). This shows that the low-contrast landmark is visible, and its presence influences their trajectories. However, the flies maintained similar speed and hover duration remained similar when approaching the combination of *low-* (Supp. Fig 3A, B) or *high-contrast landmarks* (Supp. Fig 3C, D), regardless of the distance between them.

We next pooled flight parameters for all cases in which the odorous object was low-contrast, whereas the non-odorous object was high-contrast regardless of the distance between them (blue bars, Fig 5 A-C). The distributions for these data were compared with data from cases in which the odorous object was high-contrast, but the non-odorous object was low-contrast (red bars, Fig 5 A-C). Flies flew consistently slower (Fig 5A) and hovered more (Fig 5B) for similar duration (Fig 5C) in the *low-contrast odorous landmark* treatments (blue) compared to *high-contrast odorous landmark* (red).

**Figure 5:**
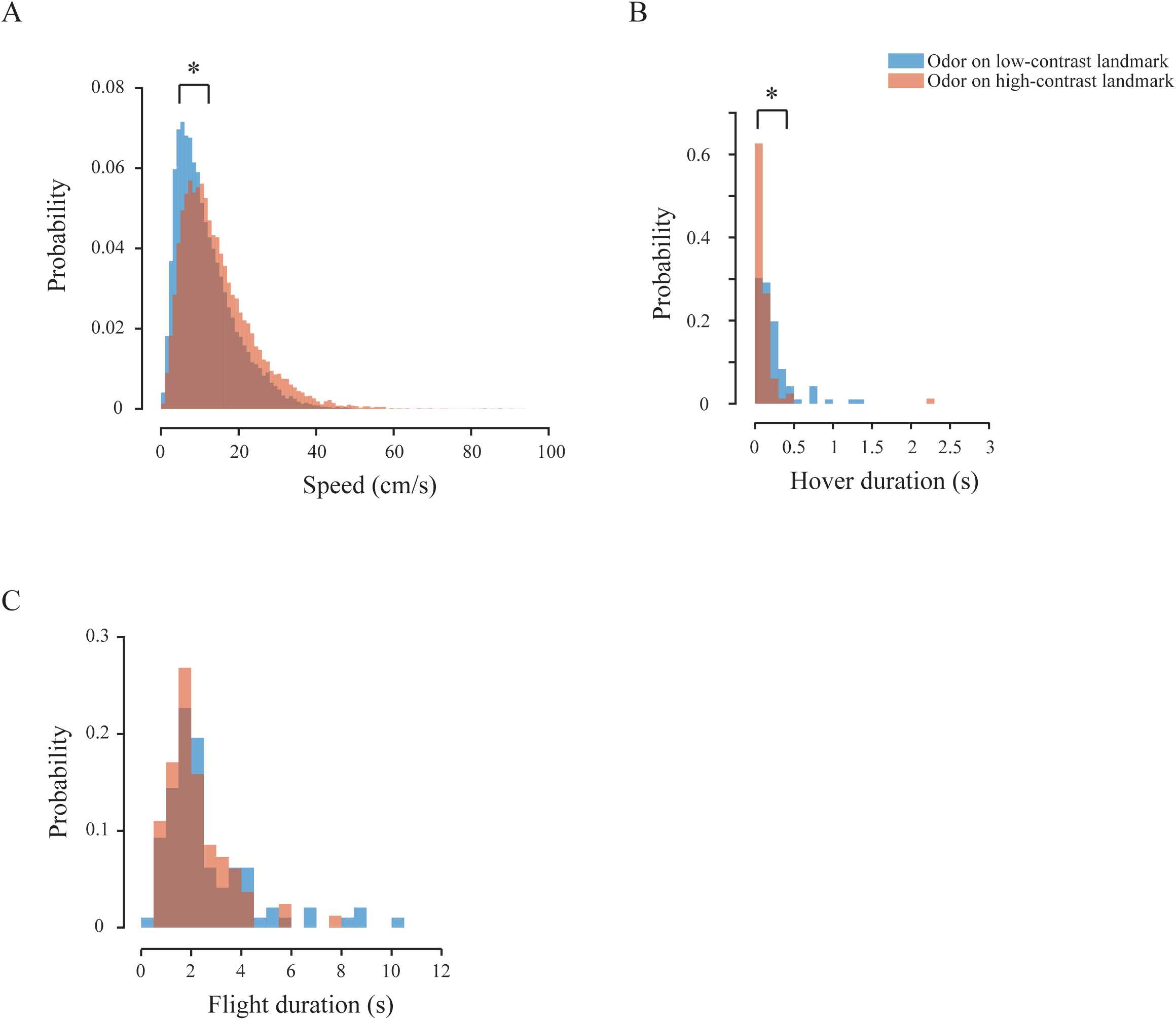
Probability distribution of flight variables from trials in which odor cues are separate from visual landmark cues. The probability distributions for speed (A), hover duration (B) and flight duration (C) in the presence of low-contrast odor source (blue) and high-contrast odor source (peach). Flight variables for treatments involving low-contrast odorous landmark were pooled (Experiment 2; N=97) and compared with the pooled variables from experiments involving high-contrast odorous landmark (Experiment 3; N=83). Frequency distribution was obtained by binning the flight variables (speed – 1 cm/s bins; hover duration – 0.1 s bins; flight duration – 0.5 s bins). Because the sample sizes in both experiments were different, we normalized the occurrences in each bin with the total occurrences for each experiment, to obtain the probability distributions from the frequencies. Statistical comparisons were conducted directly on the raw flight variables. Asterisk depicts statistically significant differences in the flight variables (p < 0.05, Kruskal Wallis test).

The above experiments suggest that flies rely on the synchrony of visual and odor stimuli to make the decision to land i.e. visual objects do not elicit landing behavior *unless* accompanied by odor cues, and *vice-versa*. Moreover, visual contrast of non-odorous objects strongly influences the landing decisions in flies, especially in the vicinity of the odor plume. This means that flies would have difficulty in finding an odor source in a visually cluttered environment, a prediction that we tested in the next set of experiments.

### Visual clutter density influences landing on odor sources

We presented flies with multiple *high-contrast landmarks,* only one of which was odorous. This created a visual clutter of several identical landmarks from which flies had to choose the correct odor source. We then tested their odor localizing ability at *low* and *high* density of visual clutter. For the *high visual clutter density* treatment, we placed seven identical landmarks at 1 cm separation from each other (Fig 6A). These were followed by two treatments with *low visual clutter density*; one with seven landmarks (Fig 6B), and another with only three landmarks (Fig 6 C) separated by 3 cm.

**Figure 6:**
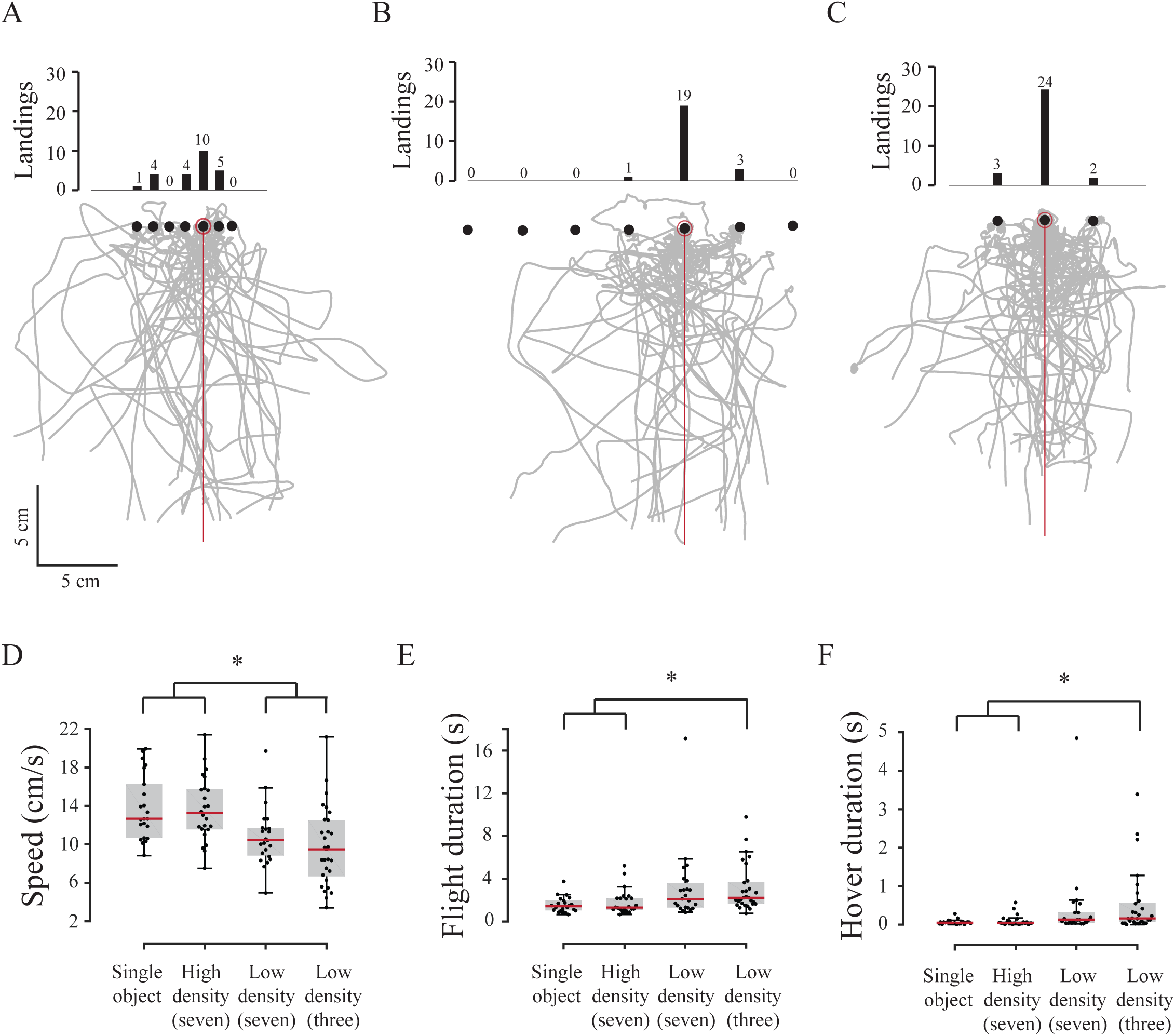
Odor tracking in visual clutter. Flight trajectories in the presence of (A) high density visual clutter (seven landmarks at 1 cm separation, N=24), (B) low density visual clutter (seven landmarks at 3 cm separation, N = 23) and (C) low density visual clutter respectively (three landmarks at 3 cm separation, N = 29), respectively. Comparison of flight parameters of three cases with a single object scenario for (D) speed, (E) flight duration and (F) hover duration.

Flies were more likely to land on non-odorous landmarks when visual clutter density was greater (Fig 6 A-C; Table 1), with a majority of the incorrect landings occurring on landmarks immediately adjacent the odor source. Increased separation between odorous and non-odorous landmarks elicited more elaborate search trajectories (Fig 6 A-C); flies flew significantly slower (Fig 6 D), increased the flight duration (Fig 6 E) and hover duration (Fig 6 F) when the visual clutter density was low. Surprisingly, flies in the high-density visual clutter flew at speeds statistically indistinguishable from a single *high-contrast odorous landmark* treatment (Fig 6 D) and their tortuosity was also not affected by the addition of multiple landmarks (Supp. Fig 3 E). Thus, presence of a low-density visual clutter meant that flies searched more and longer for the odor source, and were more likely to find the correct odorous object as compared to high-density clutter.

### Flies decrease their speed when they encounter odor

In the above experiments, flies consistently decreased their speed upon intercepting the odor plume, the approximate location of which was determined using smoke visualization (Fig 1B-C; Supp. Fig 1; see Methods and Supplementary video). How does an encounter with odor plume influence their flight on an instantaneous basis? To address this question, we examined trajectories of the flies immediately before and after the plume encounter within our region of interest. To avoid confounding the flight-related *vs* landing-related speed changes, we analyzed the data for only those flies that first encountered the plume at a distance of greater than 4 cm from the landing point (odor encounters in the range of 4-8 cm from the odor source location were analyzed). A comparison of the speeds of individual flies 250 ms before and after they intercepted the odor plume revealed that their speed decreases sharply in the time duration of approximately 50-100 ms immediately following plume encounter (Fig 7A-D; also see Supp. Fig 4 for more examples of trajectories). These speed changes are not part of their regular repertoire, as shown by the absence of changes in speed in the 250 ms duration before and after an arbitrary time point 1000 ms pre- and post-odor interception in each fly (Supp. Fig 5). However, their speed distribution shifts to lower speeds (Fig 7C) and their mean speed decreases (Fig 7D) immediately after encountering the odor plume.

**Figure 7:**
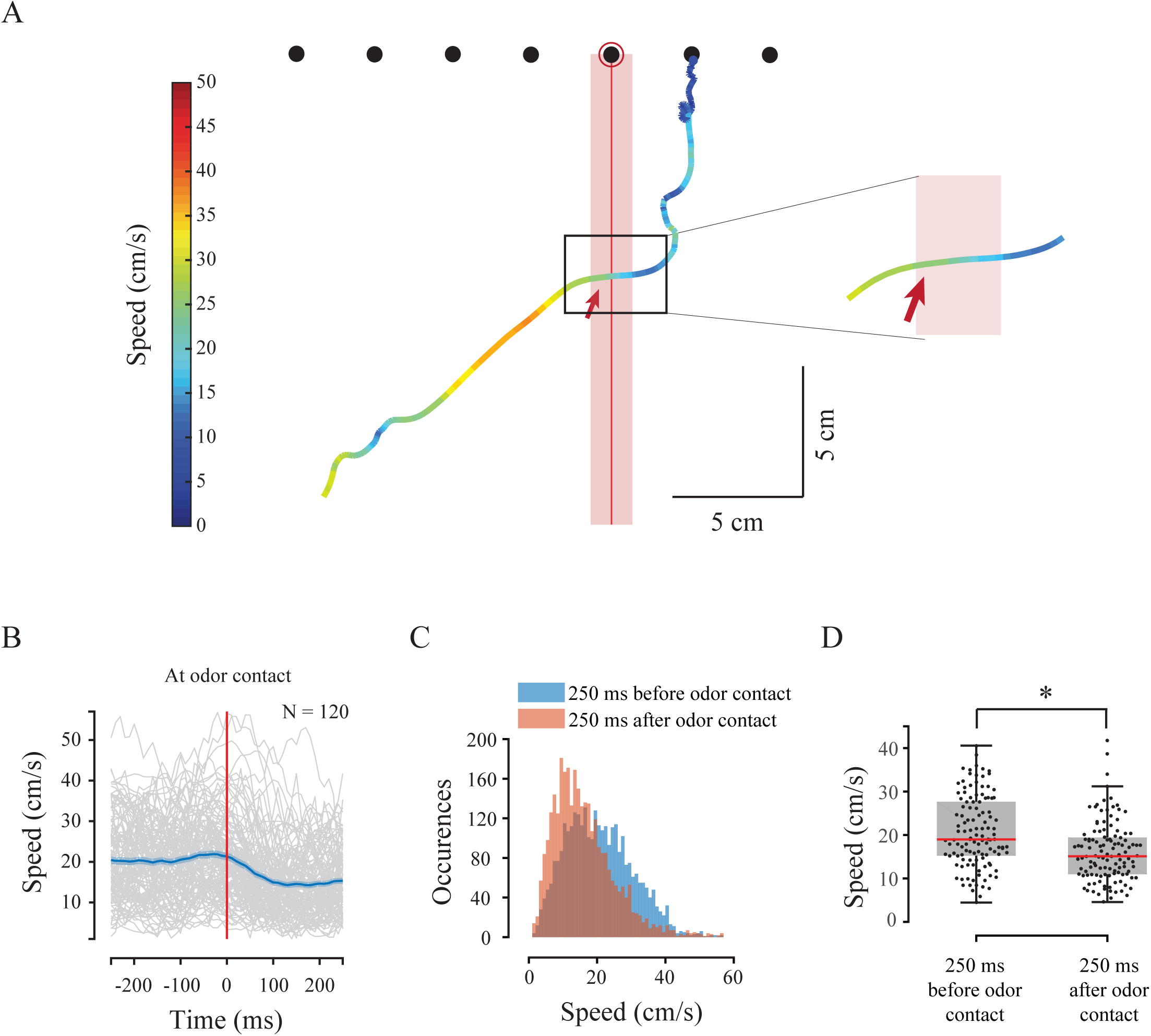
Odor encounter modulates the speed of flies. (A) A sample trajectory of a fly following odor contact with the plume (red bar of 1.6 cm width), with a closer view in the inset. Colors represent speed as indicated in the colorbar on the left, and red bar represents plume location. (B) Speed at first odor contact shown in a 500 ms window centered on the likely odor contact (250 ms before and 250 ms after odor contact). Individual speed-time curves (grey) are overlaid by the mean (blue) and standard error (light blue) (N = 83). To avoid the confounding effects of speed changes due to landing responses, only flies that encountered odor at least 4 cm before landing were used in this analysis. The decrease in flight speed is observed in less than 100 ms after first odor encounter, but not in the regions before or after the first odor encounter (also see Supplementary figure 5). (C) Speed distributions of flies upon odor contact (500 ms window). Speed distributions shifted to the lower values after odor contact. (D) Mean speeds after the first odor encounter were significantly lower than speeds before the encounter.

### Flies can localize odor sources in the absence of airflow

How do flies alter their search strategies in absence of airflow when directional cues are not clear? To address this question, we conducted trials that required flies to locate a low-contrast odorous landmark in still air. The trajectories of these flies were not directionally biased, and spread uniformly around the odor source (Fig 8A). Such flies were significantly faster (Fig 8B) and hovered less (Fig. 8C) than those in presence of airflow, however flight parameters such as flight duration and tortuosity remained unchanged (Supp. Fig 6A, B). The flight speed in still air conditions (red line; Fig 8D) was consistently greater than in flies tracking a plume in presence of the airflow (blue line, Fig 8D).

**Figure 8:**
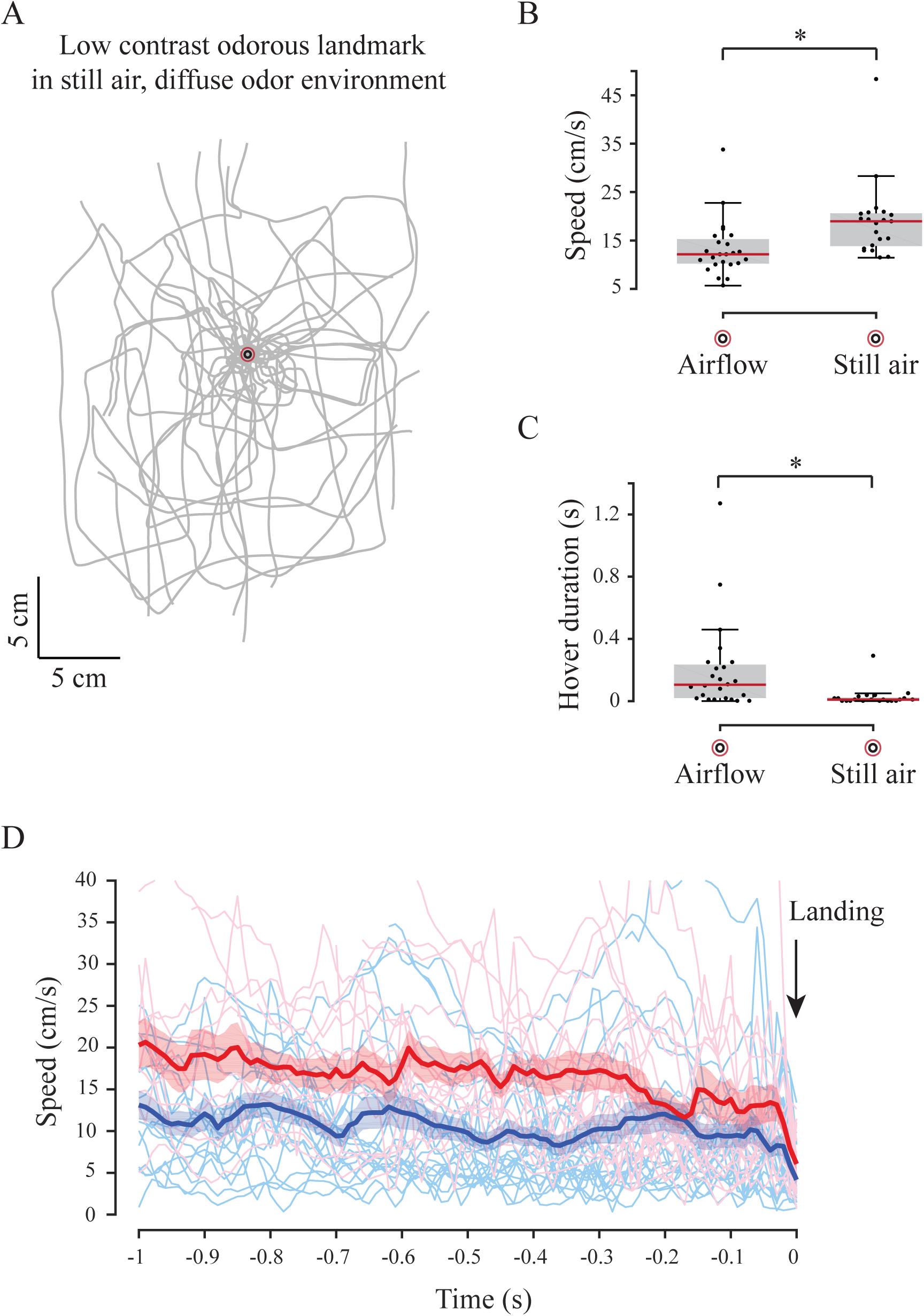
Odor tracking behavior in the absence of airflow cues. (A) Trajectories of flies in the presence of an odorous low-contrast landmark in the absence of airflow (N = 21; see Methods for details). Flies flew at significantly greater speeds (B) and hovered less (C) when airflow was absent. (D) Speed of odor tracking flies in the absence (red) *vs.* presence of airflow cue (blue) for 1s before landing. The light colored lines indicate speeds of individual flies and thick lines indicate their respective means. Shaded regions around the thick lines are the standard error of the mean.

How is the ability of flies to distinguish odorous vs. non-odorous landmarks impaired in still air? We presented the flies with two *high-contrast landmarks*, only one of which was odorous. These landmarks were separated by 1, 2 and 5 cm respectively (Fig. 9A-C). Flies performed poorly in identifying the odorous landmark when the separation between the landmarks was 1 cm (only 60% correct landings, top panel, Fig 9A), but their performance improved when the separation between odorous and non-odorous landmarks was increased (75% and 84.2% correct landings for separation of 2 and 5 cm respectively, Fig 9B,C). Flies travelled for longer durations (Fig 9D) with greater tortuosity (Fig 9E) in the 2 cm separation case as compared to 5 cm. However, their speed and hover duration were not significantly different for any arrangement of these objects (Supp. Fig 6C, D).

**Figure 9:**
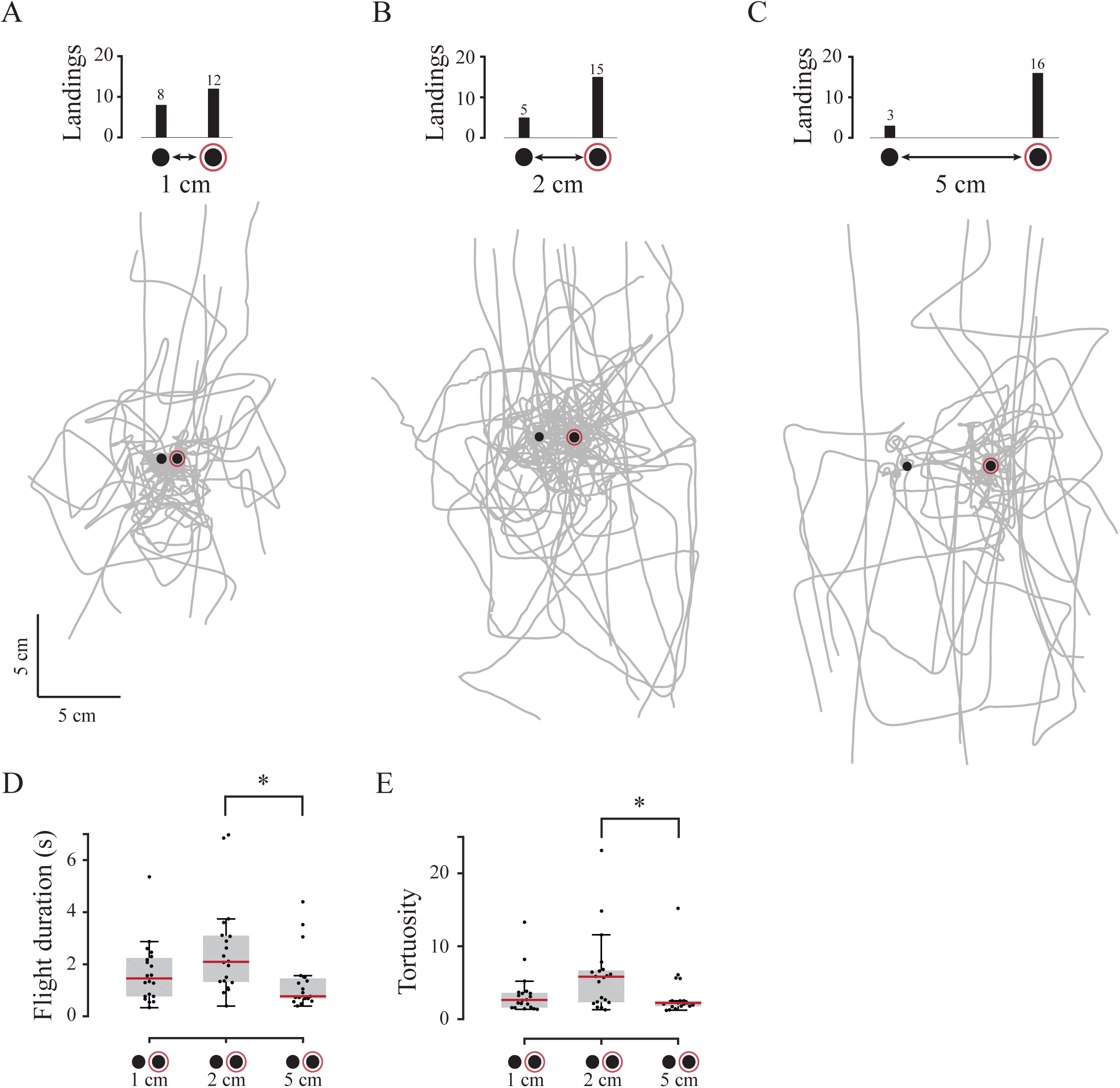
Flight trajectories in the presence of a high-contrast odorous landmark paired with an identical non-odorous landmark in absence of airflow. These landmarks are separated by (A) 1 cm (N = 20), (B) 2 cm (N = 20) and (C) 5 cm (N = 19) respectively (see Methods for details). Bar plots in the upper panel show the number of landings on each object. A comparison of the flight durations (D) and tortuosity (E) are shown as box-and-whisker plots.

In still air, when flies had to find a *low-contrast odorous landmark* separated from a *high-contrast non-odorous landmark* by 5 cm, they landed on both objects with roughly equal probability (57% correct landings; Table 1; Fig 10A). On the other hand, they could very reliably find a *high-contrast odorous landmark* when it was separated by a *low-contrast non-odorous landmark* in still air (95% correct landings; Fig 10B). None of the flight parameters in these two cases were significantly different from each other (Fig 10 C-F). Thus, their choice is substantially biased toward high-contrast visual objects in absence of a plume to guide them. It is also illustrative to compare these treatments with those in which airflow was present for similar object arrangement (Fig 3C, Fig 4 C) for which airflow and odor plume were present. The ability of the fly to find the odor source was substantially enhanced by the presence of the plume, which underscores its importance in odor tracking behavior. Note also that in both moving and still air, flies tended to hover in front of objects just before landing (compare Supp. Fig. 8 A, B with Fig 8 C, D).

**Figure 10:**
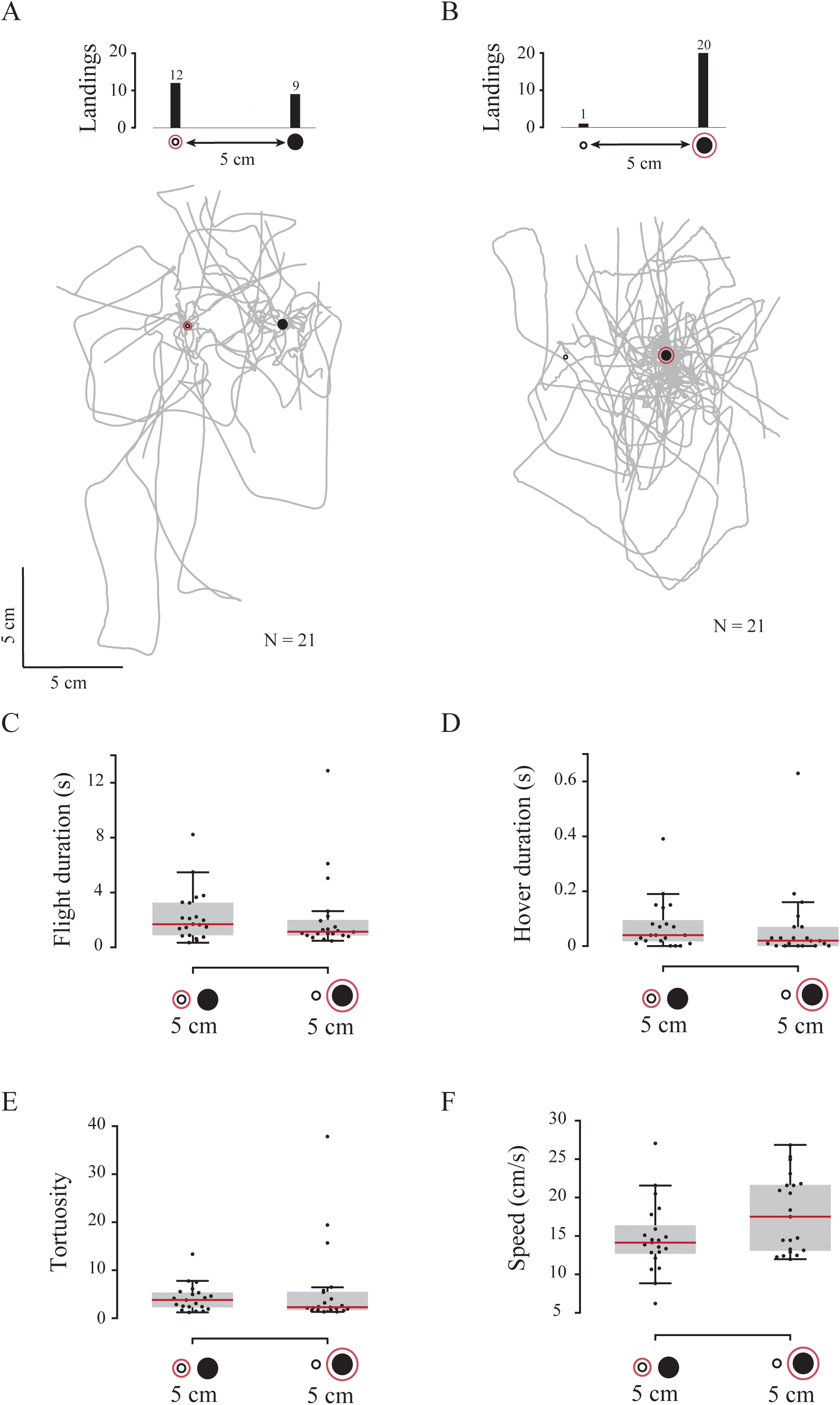
Landing preferences for low-contrast vs. high-contrast landmarks in the absence of airflow. Flight trajectories (grey lines) for (A) an odorous low-contrast landmark presented in combination with a non-odorous high-contrast landmark (N = 21) in contrast to (B) an odorous high-contrast landmark presented with a non-odorous low contrast landmark (N = 21), separated by 5 cm in both the treatments. The comparisons for (C) flight duration, (D) hover duration, (E) tortuosity and (F) speed between the above two treatments revealed no statistical differences (Kruskal Wallis test at 95% level of significance).

Thus, synchronous odor and visual cues are also essential for odor source location in still air conditions. Barring localized micro-flows (e.g. due to wing motion), the odor in this case spreads largely through diffusion, to form a gradient, which the flies appear to track in still air.

## Discussion

Locating an odor source in a visually-cluttered environment is a complex task which requires inputs from multiple senses, including the visual and olfactory modalities which then drive motor responses (e.g. Raguso and Willis, 2002; Frye et al., 2003; Dekker and Cardé, 2005). For flying insects, this means controlling flight in three dimensions in environments that are typically turbulent (Murlis, 1992). Because the proper identification of odor sources is essential to gain access to food and mates, the question of how insects solve this problem has been of central importance to biologists over several decades (e.g. Kennedy, 1983; Raguso and Willis, 2002).

What basic rules guide the flies to odor sources in visually ambiguous conditions? Previous studies have outlined several specific behaviors including optomotor anemotaxis, cast-and-surge maneuvers, odor-guided salience changes etc. which enable insects to arrive in the vicinity of an odor source (e.g. Kennedy and Marsh, 1974; Vickers, 2000; Chow and Frye, 2008). Our study sought to specify how insects, having arrived in the general region of an odor source, pinpoint its precise location from among many possibilities in the decisive moments before landing.

### Odor resolution is vision-dependent

One key finding of this study is that when flies encounter an odor plume that indicates the presence of a potential food source, they *decrease* their speed with a latency of under 100 ms (Fig. 7A-D). This behavior may serve two functions: first, it provides the flies with greater sampling time to determine the spatio-temporal co-occurrence of odor and visual feedback. Second, it increases the probability of repeated odor encounters, which would enable flies to determine the general orientation of an odor source. These observations contrast with previous studies which showed that flies *increase* their groundspeed approximately 190 ± 75 ms following a plume encounter (Budick and Dickinson, 2006; van Breugel and Dickinson, 2014; Bhandawat et al, 2010). However, in those studies there were no visible landmarks at the time of odor encounter, and hence landing was not imminent. In contrast, the trajectories reported here were derived from a region that was between 4-8 cm from the nearest visible odor source. This suggests that odor encounter triggers a behavioral switch in flies that causes them to seek visual objects, even though these had no inherent salience when odor was absent (Fig. 2A; also Budick and Dickinson, 2006). We also show an increased bias towards objects of a higher visual contrast and situated in the immediate vicinity of the odor source (Fig. 3A-C, 6 A-C), which is consistent with van Bruegel and Dickinson (2014). The bias towards high-contrast objects means that flies may sometimes incorrectly identify the odor source location if it does not exactly overlap with a visual landmark (Fig 3A). However, when the two objects are sufficiently separated, flies are more successful at correctly identifying the odor source location (Fig. 3C). Thus, flies depend on the spatiotemporal co-occurrence of visual and odor cues to identify the odor source, and their odor resolution is vision-dependent.

In presence of multiple landmarks (visual clutter), flies initiate a search behavior which is characterized by slower speed, increased tortuosity and longer flight / hover duration (Fig. 3D-E, 6D-F). This may help ascertain the co-occurrence of visual and odor cues by allowing for more time to process odor. The limited resolution of their compound eyes means that flies may not correctly pinpoint the odor source location within a high-density clutter (Fig. 6A). Their search behavior is significantly enhanced when the location of the landmark does not match with odor cue. In contrast, a single odorous landmark does not elicit an elaborate spatial search. Instead, flies steadily decrease their distance from the odor plume axis while approaching the target thus honing in on the odor plume, regardless of whether the landmark was high-(Supp. Fig 2C) or low-contrast (Supp. Fig 2D). These findings demonstrate the dominant influence of visual landmarks during odor searches, which are especially important in natural scenarios.

### Flies use a different strategy for odor tracking in absence of airflow

How do insects find odor sources in still air conditions? Although airflow is an important cue for odor-seeking insects (e.g. Kennedy and Marsh, 1974 Budick and Dickinson, 2006; Willis and Arbas, 1991), flies could also successfully track down an odor source in still air (Fig 8-10). In static air, odor propagation is isotropic and generates uniform concentration gradients around the odor source, although these gradients may be locally disturbed by self-induced flow from flapping wings (Sane and Jacobson, 2006), possibly aiding odor detection (Loudon and Koehl, 2000). Do flies use similar strategies when tracking odor in still air? Without airflow to break the odor symmetry, flies approach the odor source equally from all directions (Fig 8A). They fly at faster speeds (Fig 8B) and hover less (Fig 8C) as they steadily hone in on the odor source, as also reported in mosquitoes tracking CO_2_ in still air (Cardé and Lacey, 2012; Breugel and Dickinson, 2015). This alternate strategy is robust because it still allows a majority of flies to find the correct odor source from two visually identical objects separated by 2 cm or more (Fig 9A-C, Table 1). Thus, flies adopt different strategies when searching the odor sources in the presence vs. absence of airflow (summarized in Fig 11).

**Figure 11:**
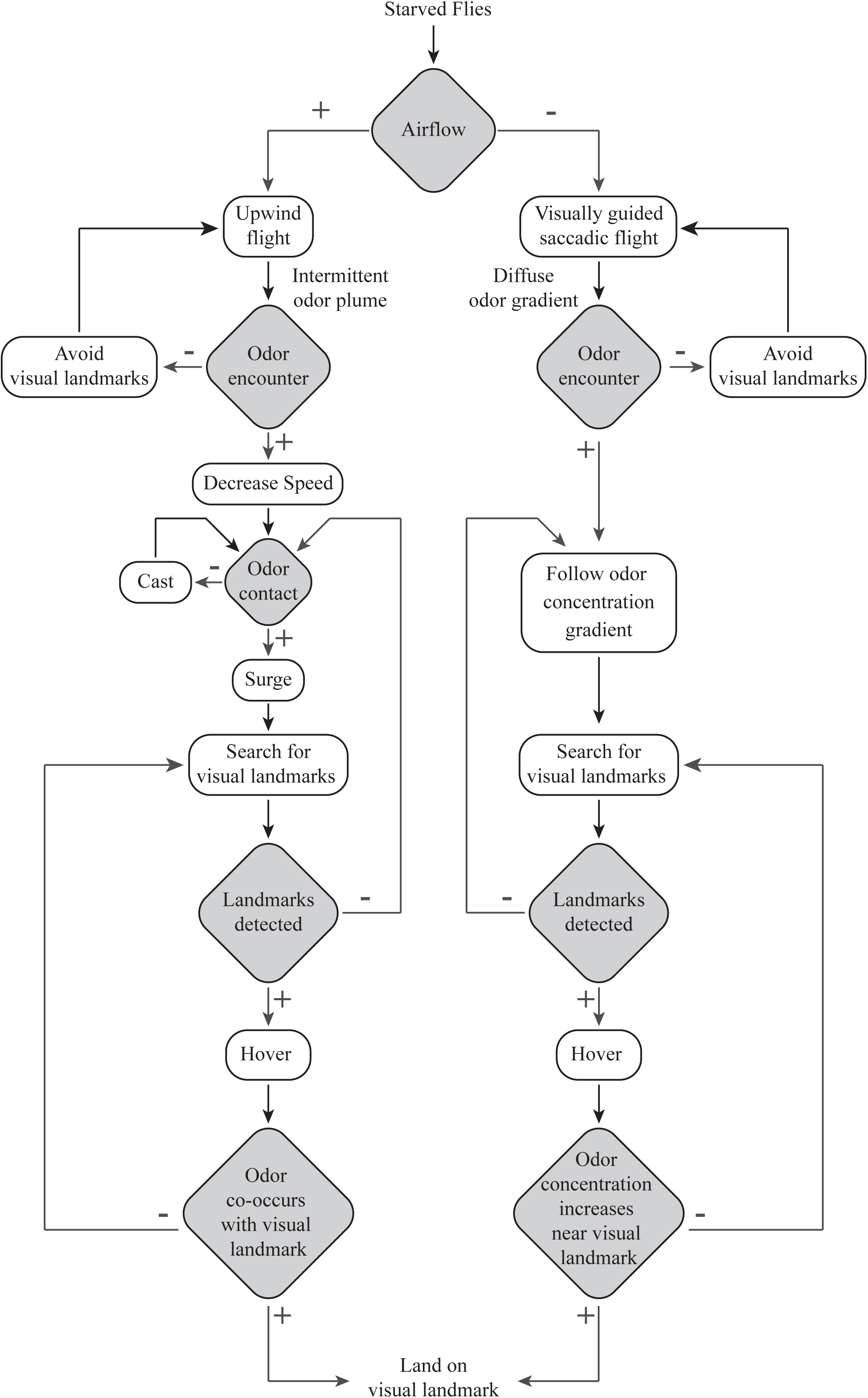
A flowchart of odor-tracking strategies in flies. A flowchart derived from previous studies and the results described here shows distinct strategies employed by flies based on presence (left) or absence (right) of airflow. In the former case, flies track plumes whereas in the latter case, they track odor gradients. + signifies the presence and – the absence of the associated cue. Grey diamond-shaped boxes display sensory cues which include airflow, odor, landmarks and their combination, and motor actions are displayed in unfilled rounded rectangles. Both strategies terminate after flies land on the landmark.

### Olfactory working memory in flies?

Whether in presence or absence of airflow, a large majority (76%) of the 311 flies tracking multiple visual landmarks across 13 different treatments landed successfully on the odor source, underscoring the robustness of combined strategies of plume- and gradient-tracking. A key ingredient of these strategies, not directly addressed in our experiments, is a neural mechanism to ensure that flies continue to search the plume even after leaving the plume. Some examples of such trajectories, for both successful and unsuccessful searches, are shown in Supp. Fig. 7.

For spatial navigation tasks, the existence of a spatial working memory has been well-demonstrated in the case of visual tracking, in which *Drosophila* flies moving between two vertical poles maintain their direction for several seconds after these landmarks became extinct or reappeared elsewhere (Neuser et al., 2008). We hypothesize the presence of an ‘olfactory working memory’, which keeps track of the previous odor encounters, and which may ensure that flies continue their search for odor sources even when odor cues become temporarily extinct. A fundamental requirement for odor working memory is to successfully register an odor encounter, and display behavior that suggests that it recalls this odor encounter. As shown in this study, flies sharply decrease their flight speed after a putative odor encounter (Fig 7 A-D). Moreover, a majority of the flies maintain an attraction towards visual landmarks even without frequent odor encounters. In the absence of airflow, a large fraction of the flies (30%, 2 cm separation; Fig 9B) iteratively approached the identical landmarks before landing on the correct odor source (Supp. Fig 9). Together, these observations suggest the possibility of an ‘olfactory working memory’, which enables them to recall a prior plume encounters for several seconds after leaving it. Future studies must quantify the duration for which this memory lasts, and where in the brain it resides.

### Visual and olfactory specialization in insects

From an evolutionary perspective, how do certain insects evolve to specialize on specific fruits or plants in their natural surroundings? Examples of such specialists have been reported in *Drosophila*, including *D. sechellia*, which specializes on the fruit, *Morinda citrifolia* (Higa and Fuyama, 1993; Jones, 2005), which is toxic to related Drosophila species but not to *D. sechellia.* Similarly, *D. pachea* are found on the rotting stems of the cactus *Lophocerus schottii* (Heed and Kircher, 1965). The bias for high-contrast visual cues *vis-a-vis* odor cues suggests the testable hypothesis that specialist insects are attracted to specific olfactory and visual cues. Such preferences have been demonstrated, for instance, in the Tephritid fly, *Rhagolettis pomonella* (Walsh) for apple-like stimuli (e.g. Aluja and Prokopy, 1993). Here, an attractive odor stimulus makes specific landmarks in the surrounding environment attractive, which in turn biases their landing decisions (Fig 2B, C). If flies or other insects have evolved to specialize on odor objects of specific visual signatures, then we expect to see strong bias towards objects of specific shape or color, or else they may be more biased towards specific odor stimuli irrespective of their visual appearance. By enabling us to demonstrate that flies make a weighted decision between odor and visual stimuli, our study thus provides the methodology to test this hypothesis.

## Conclusion

Our paper shows that during plume tracking, *Drosophila melanogaster* use both olfactory and visual cues. In the final phase of odor localization just before landing, the flies decelerate following an odor plume encounter, and they undergo a behavioral ‘switch’ that enhance salience towards high-contrast visual objects in the immediate vicinity of the odor plume. This ‘switch’ ensures that flies continue seeking the odor source even after losing direct contact with the odor. If the visual objects are far from the odor plume, flies are attracted to them but less likely to land on them. Thus, when tracking an odor plume, flies determine the presence of an odor source based on the synchrony of visual and odor cues. In still air, flies adopt a different strategy, which may involve flying down an olfactory gradient towards visual landmarks. These two strategies provide a robust means for the fly to precisely locate an odor source.

## Supplementary Figure Legends

**Supplementary Figure 1:**
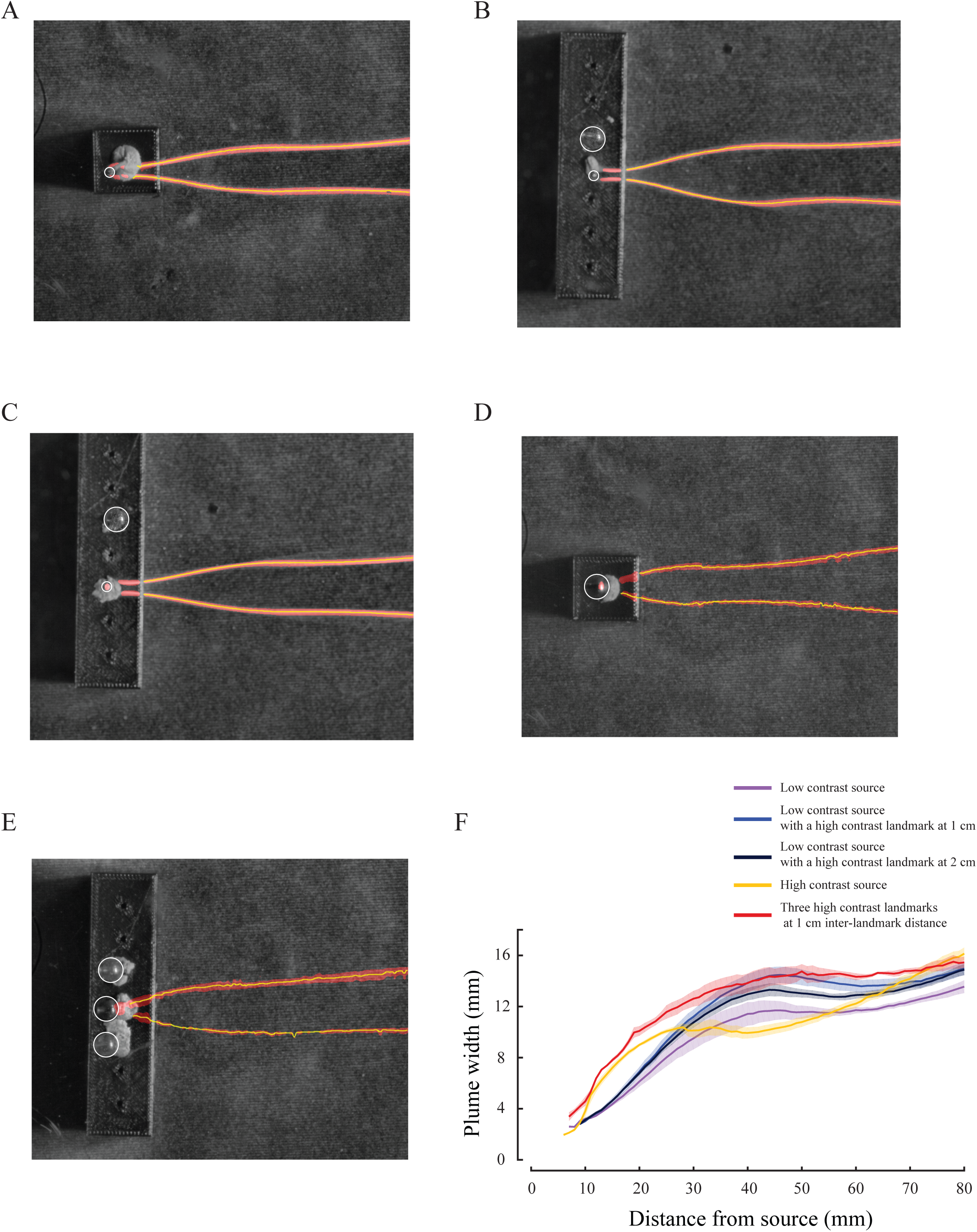
Plume visualization and quantification of plume width. Steady state smoke plume, viewed from above, for (A) Capillary (low-contrast landmark, N = 4); (B, C) Capillary (low-contrast landmark) source with spherical bead (high-contrast landmark) separated by 1, 2 cm respectively (N = 4 for both); (D) Spherical bead (high-contrast landmark, N = 4); (E) 3 spherical beads separated by 1 cm (Visual clutter, N = 4). The red bands show the averaged plume at steady-state with yellow lines indicating the median (see methods). (F) Variation in plume width vs. distance from the source along the plume axis for smoke-visualized plumes. Colors represent specific treatments. Dark lines show the mean plume width and the light bands show the standard error around mean.

**Supplementary Figure 2:**
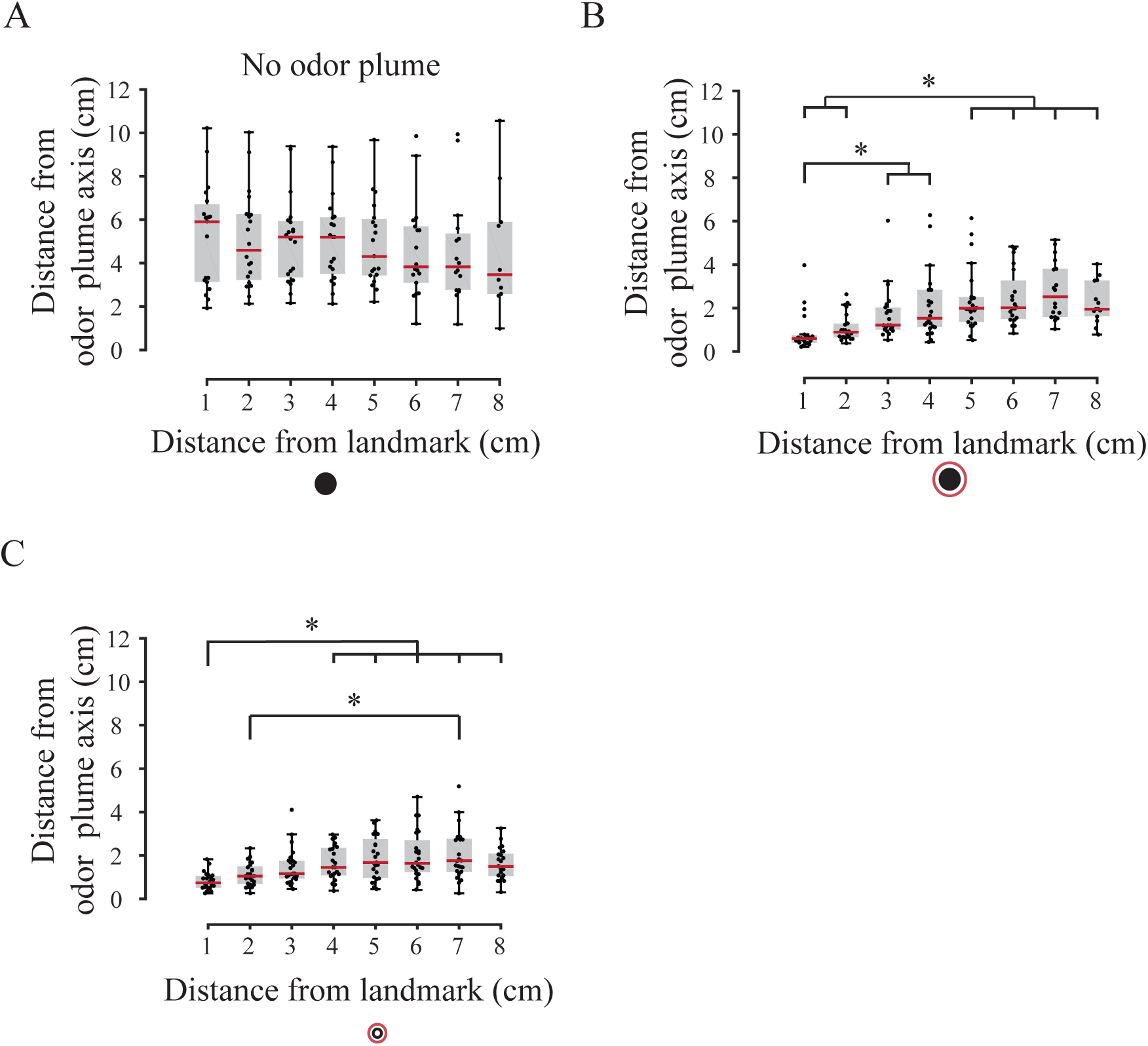
Approach behavior in the presence of non-odorous vs. odorous landmarks. A) In the absence of odor cue, flight trajectories are not along the axis of the visual object, suggesting that they do not fly towards a visual landmark along its axis (N = 20). B) In the presence of an odor cue, flies gradually decreased their average distance from the odor plume axis as they approached both a high-contrast odorous landmark (N = 22) and C) low-contrast odorous landmark (N = 24).

**Supplementary Figure 3:**
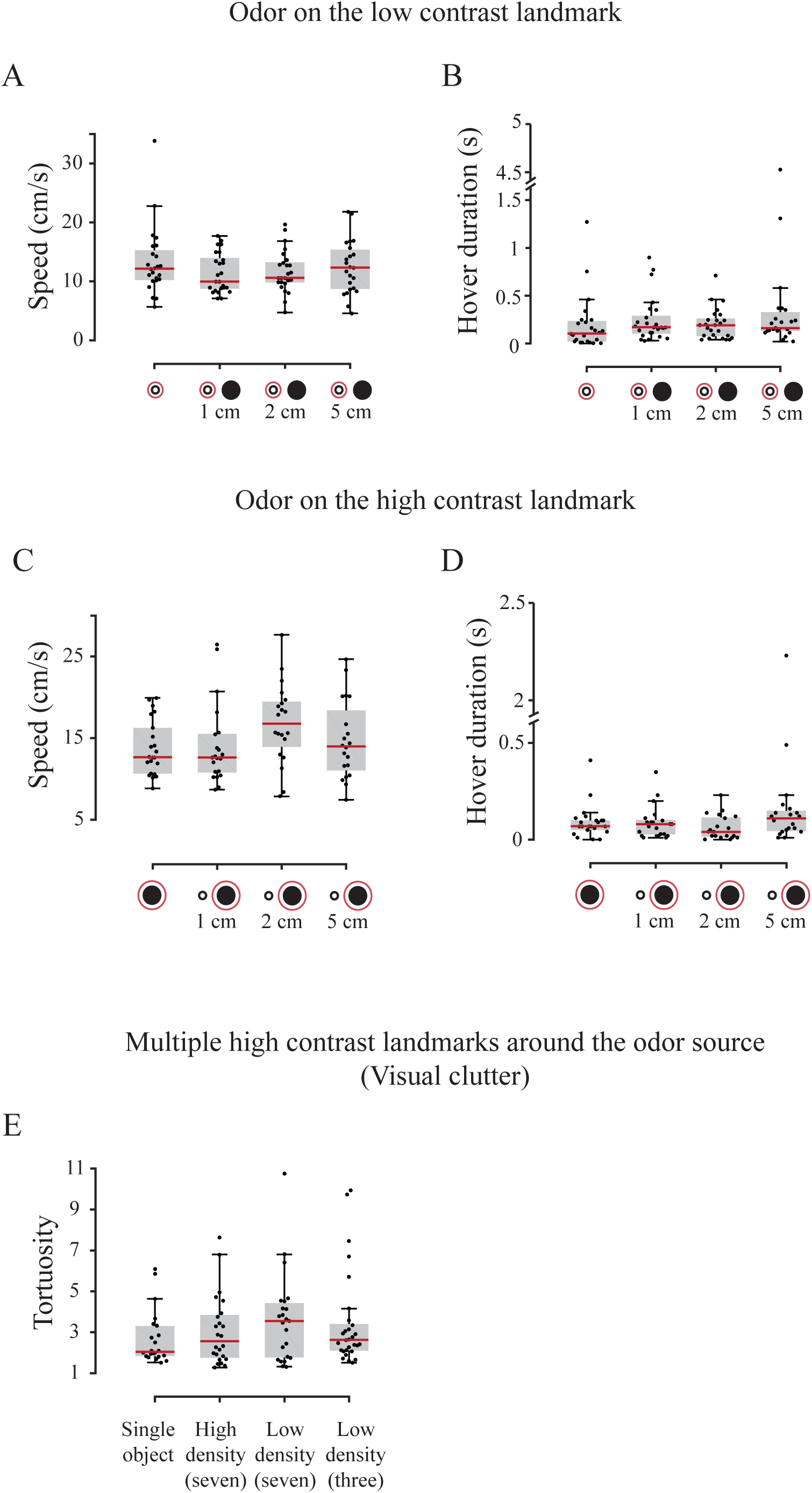
Comparison of additional flight variables for vision-odor separation and visual clutter experiments. Box-plots of flight variables for flight trials, which do not show any statistically significant differences while plume tracking in presence of visual landmarks. (A, B) low-contrast odorous landmarks are paired with high-contrast non-odorous landmarks at different separation distances (only low-contrast odorous landmark, 1cm, 2 cm and 5 cm). These include speed (A) and hover duration (B). Similar plots for (C, D) high-contrast odorous landmarks paired with low-contrast non-odorous landmarks (only high-contrast odorous landmark, 1cm, 2 cm and 5 cm) including speed (C) and hover duration (D). (E) In the visual clutter treatment, tortuosity of the flight trajectories did not change significantly across different arrangements of visual clutter densities. Details of treatments, sample sizes and statistics are provided in the Methods section of main text.

**Supplementary Figure 4:**
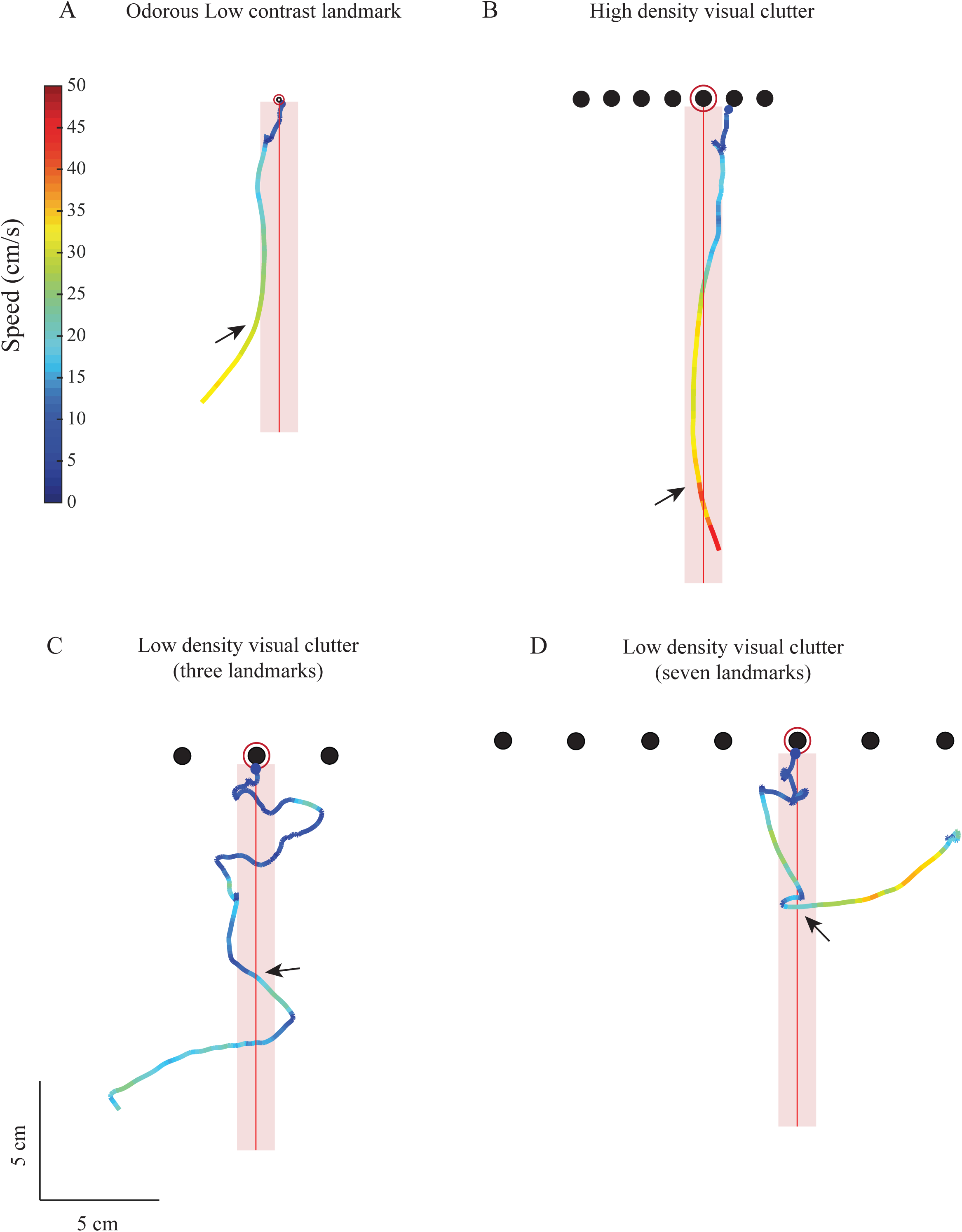
Additional examples of trajectory plots illustrating speed change in flies following a putative odor encounter. Sample flight trajectories of flies flying in the presence of different landmark arrangements. These trajectories are examples of how instantaneous speed of flies decelerates after odor encounters (arrows). Odor plume axis (red line) is surrounded by the cylindrical odor plume, assumed to be approximately 1.6 cm wide (light red band). Flight speed is depicted using a color map. Shown here are sample flight trajectories in the presence of (A) a single low-contrast odorous landmark, and a combination of low-contrast odorous landmark and a high-contrast non-odorous landmark at (B) 1 cm and (C) 2 cm separation. (D) Sample trajectory in the presence of low-density visual clutter of 3 landmarks with odor on the central landmark.

**Supplementary Figure 5:**
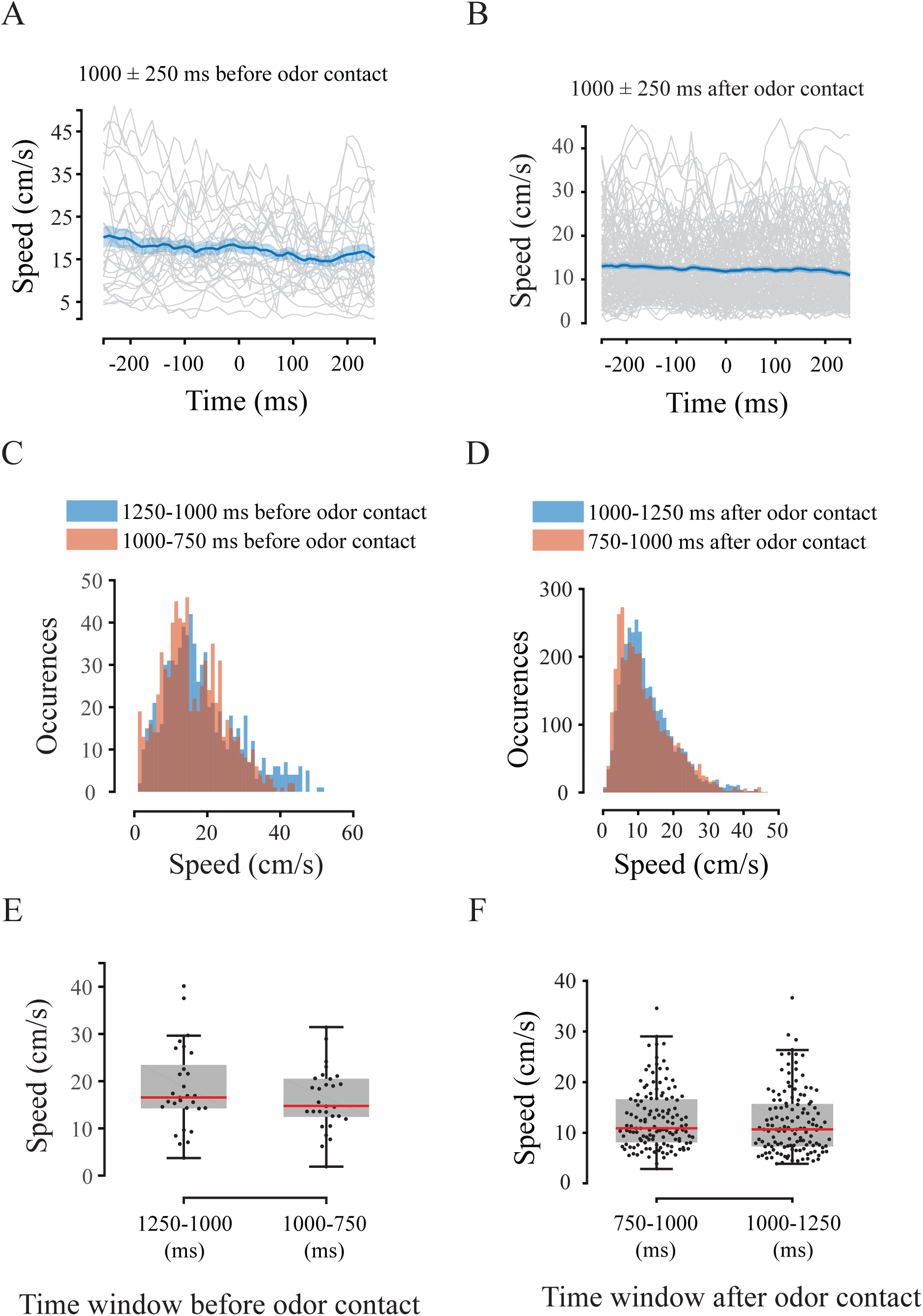
Speed change before and after odor encounter. (A, B) Speed vs. time for individual flies 1250 ms before (N = 29) and after (N = 134) first odor contact (grey; 500 ms window). The mean (blue) and the standard error of the individual speeds (light blue) of these plots are also shown. Only flies that first encountered odor at least 4 cm before landing were used for this analysis to avoid landing related speed changes (see Methods). (C) Speed distributions of flies 1250-1000 ms (peach) and 1000-750 ms (blue) before odor contact (500 ms window). (D) Speed distributions of flies 1250-1000 ms (peach) and 1000-750 ms (blue) after odor contact (500 ms window). Distributions remain similar during both pre and post time windows. (E, F) Speed values were not significantly different during both pre and post 1250ms of odor encounter. (p<0.05, Kruskal Wallis test, Nemenyi's test).

**Supplementary figure 6:**
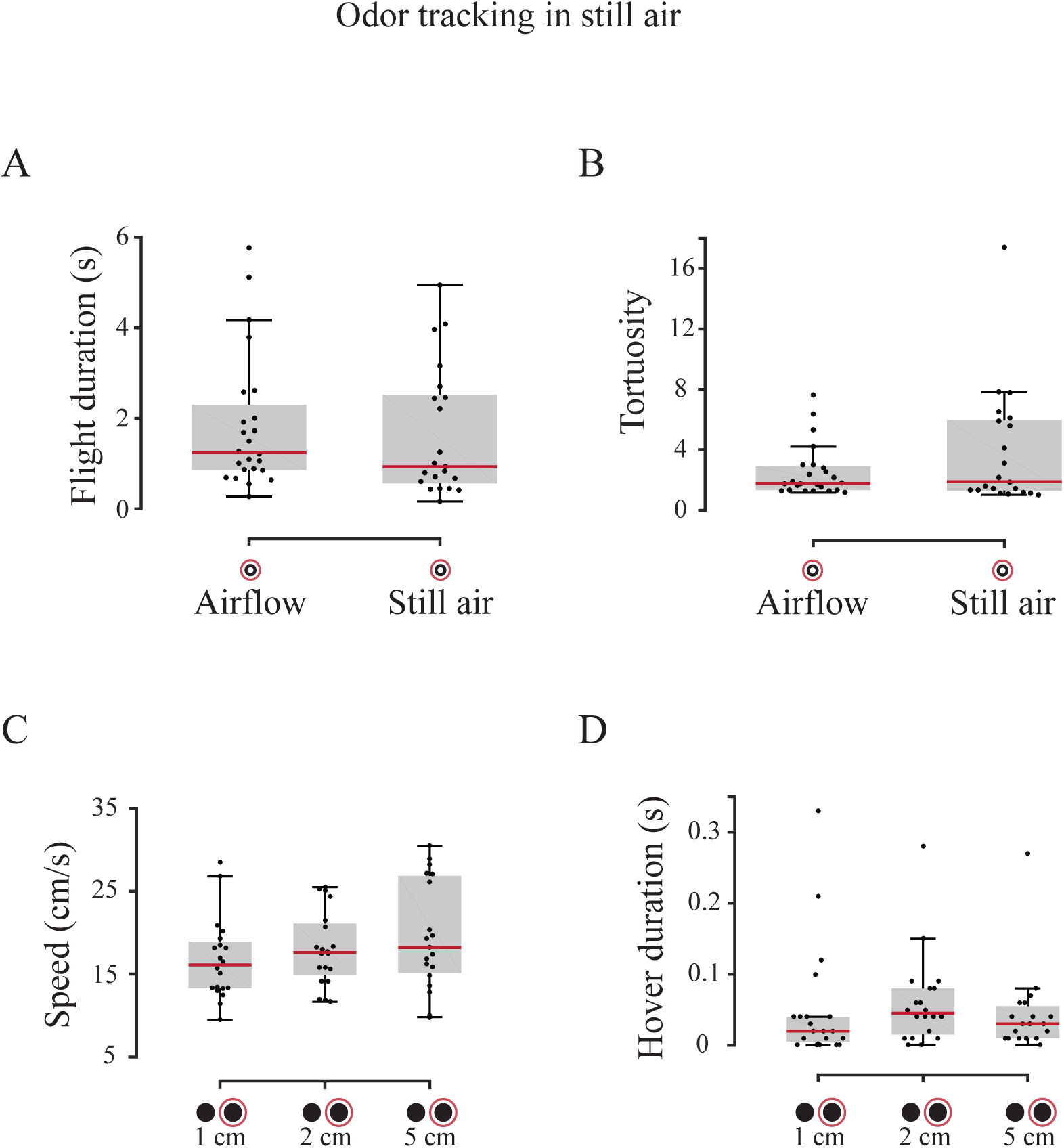
Box-plots of flight variables for flight trials, which show no statistically significant differences in presence of still air and visual landmarks. A) Flight duration and B) tortuosity of the flies tracking a low-contrast odorous landmark did not significantly vary with the presence or absence of airflow cue. Flies also did not have significant differences in the C) speed and D) hover duration, when tracking high-contrast odorous landmark in the presence of a high-contrast non-odorous landmark at various separation distances (1, 2 and 5 cm).

**Supplementary figure 7:**
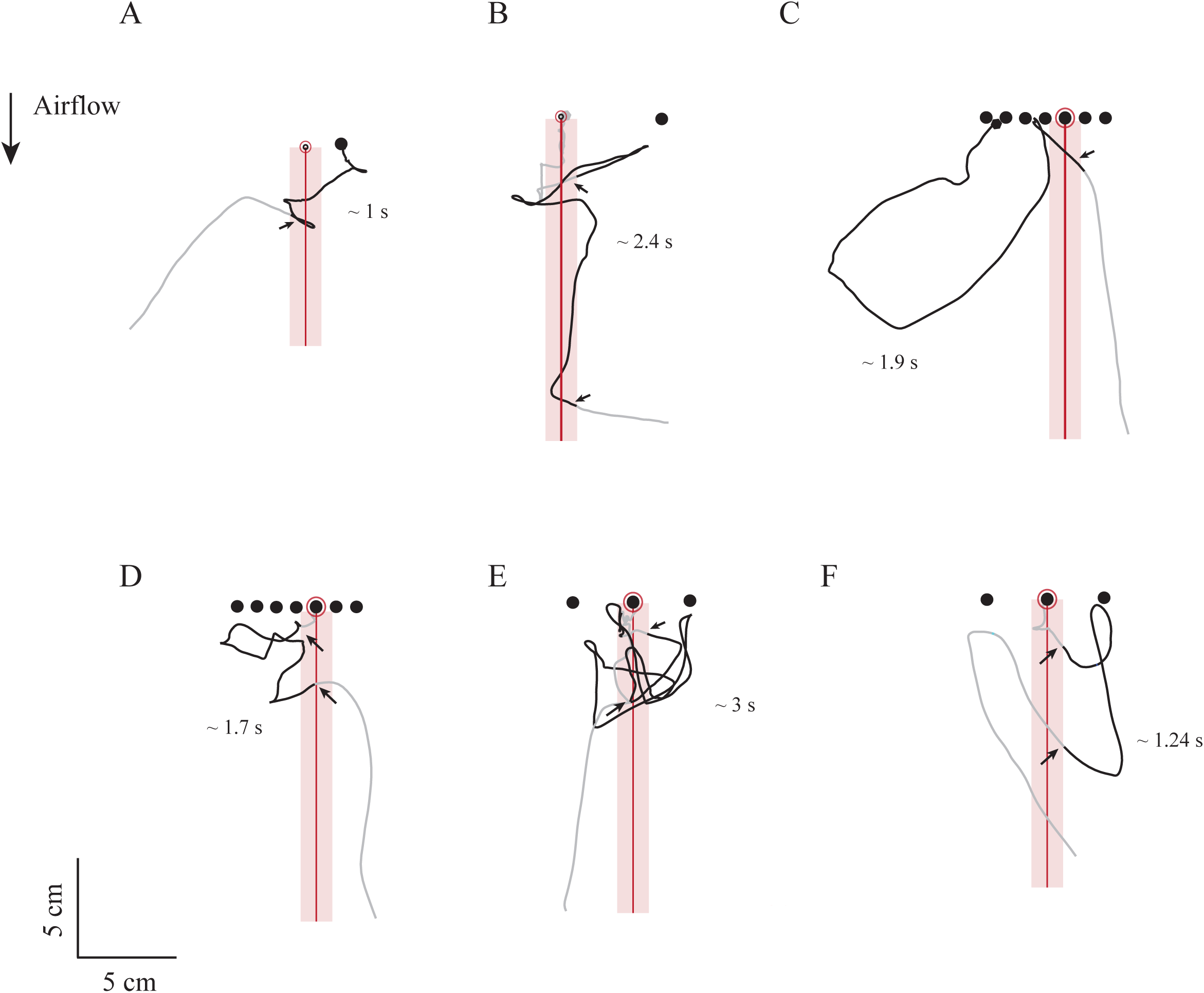
Sample flight trajectories of flies that sample odorous landmarks after leaving the odor plume. Sample flight trajectories in which flies are away from the plume (of approximate width of 1.6 cm, see methods) for approximately 1 sec (A), 2.4 sec (B), 1.9 sec (C), 1.7 sec (D), 3 sec (E), and 1.24 sec (F). The segments of flight trajectories in which flies were outside the odor plume after odor contact are highlighted in black and the rest of the trajectory is shown in gray color. (F) The sample flight trajectories obtained from treatments in Experiment 2 (A, B) and from Experiment 4 (C-F). Such flight trajectories suggest that flies can maintain odor tracking behavior and landing even without immediate odor encounters.

**Supplementary Figure 8:**
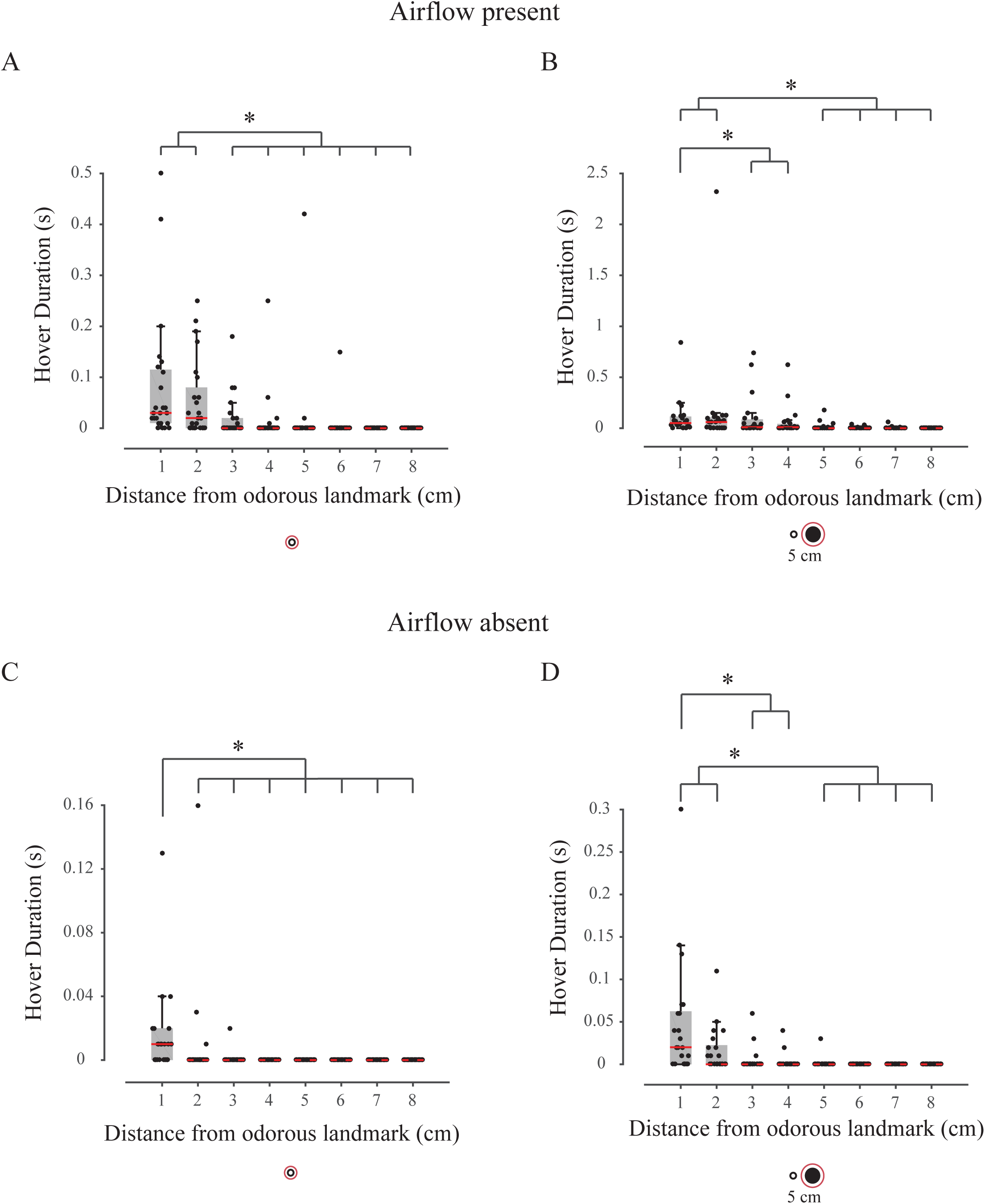
Hover duration vs. distance from the odor source in presence and absence of airflow. Total hover duration in the presence of airflow for A) low-contrast odorous landmark and B) high-contrast odorous landmark and low-contrast non-odorous landmark at 5 cm separation. Similarly, the total hover duration in the absence of airflow for C) low-contrast odorous landmark and D) high-contrast odorous landmark and low-contrast non-odorous landmark at 5 cm separation. Hover duration prior to landing increases both in the presence and absence of airflow. Statistically significant differences are indicated by the asterisk symbol above the box-plots (p<0.05, Kruskal Wallis test, Nemenyi's test; see Methods for details).

**Supplementary figure 9:**
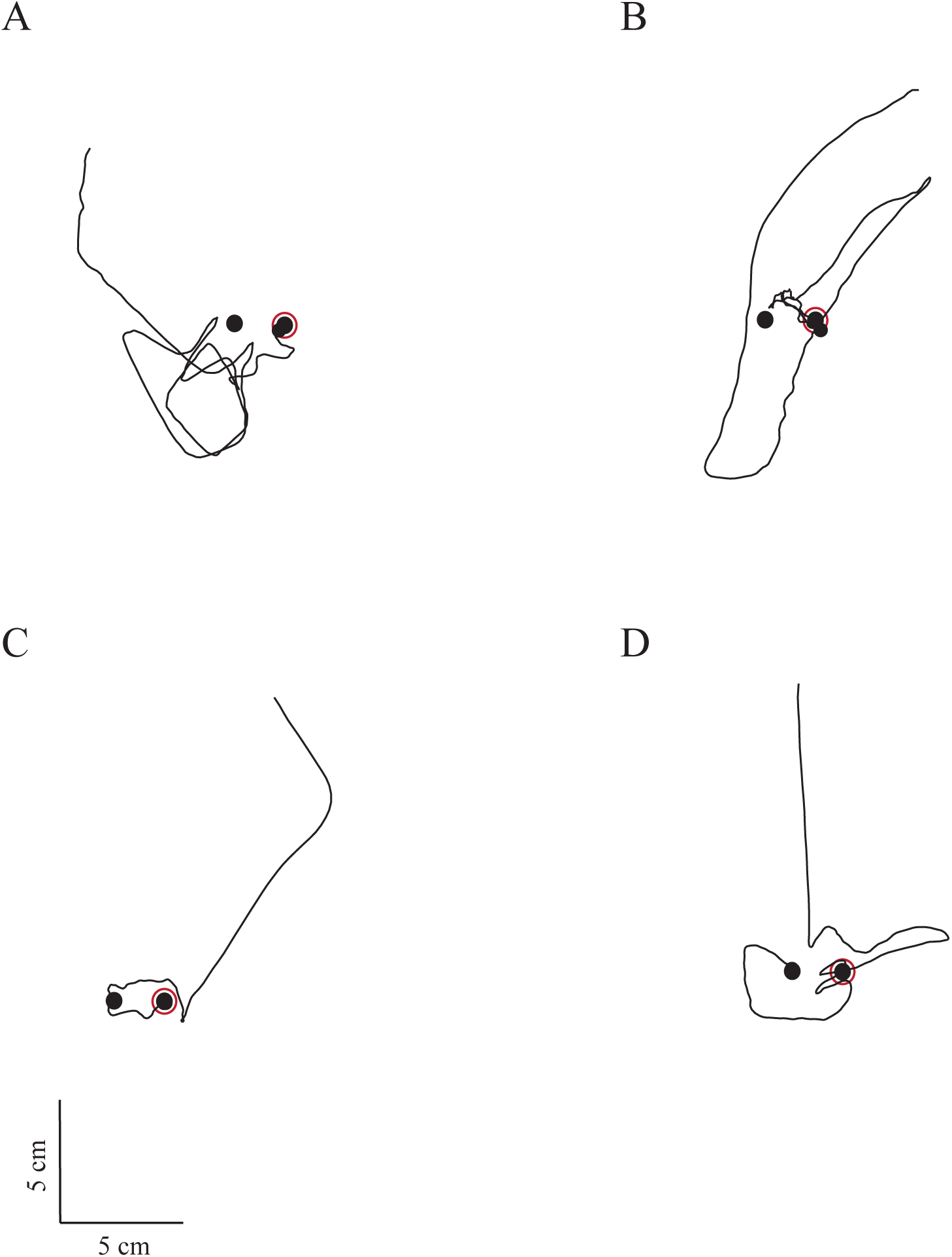
Sample trajectories illustrating odor-tracking in absence of airflow. Examples of flight trajectories from a treatment in which the separation between the odorous and non-odorous landmark was 2 cm in absence of airflow (Experiment 6). (A-C) Examples in which flies found the correct location of the odorous landmark after search and (D) example in which the fly landed incorrectly on the non-odorous landmark despite search. These examples are presented to highlight both the robustness and difficulty inherent in searching for an odor source in absence of airflow.

